# The repertoire of copy number alteration signatures in human cancer

**DOI:** 10.1101/2022.11.14.516412

**Authors:** Ziyu Tao, Shixiang Wang, Chenxu Wu, Tao Wu, Xiangyu Zhao, Wei Ning, Guangshuai Wang, Jinyu Wang, Jing Chen, Kaixuan Diao, Fuxiang Chen, Xue-Song Liu

**Author notes:** These authors contributed equally to this work (Ziyu Tao, Shixiang Wang, Chenxu Wu).

## Abstract

Copy number alterations (CNAs) are a predominant source of genetic alterations in human cancer and play an important role in cancer progression. However comprehensive understanding of the mutational processes and signatures of CNA is still lacking. Here we developed a mechanism-agnostic method to categorize CNA based on various fragment properties, which reflect the consequences of mutagenic processes and can be extracted from different types of data, including whole genome sequencing (WGS) and SNP array. The 14 signatures of CNA have been extracted from 2778 pan-cancer analysis of whole genomes (PCAWG) WGS samples, and further validated with 10851 the cancer genome atlas (TCGA) SNP array dataset. Novel patterns of CNA have been revealed through this study. The activities of some CNA signatures consistently predict cancer patients’ prognosis. This study provides a repertoire for understanding the signatures of CNA in cancer, with potential implications for cancer prognosis, evolution, and etiology.

## Background

Somatic mutations are the driving force of cancer development. Genomic alterations in cancer cells consist of two major categories: (1) small scale alterations that include single base substitutions (SBSs) and small insertion and deletions (INDELs), and (2) large scale alterations known as structural variations. Copy number alteration (CNA) is a major type of structural variations, and is prevalent in human cancer [1, 2]. CNAs and SBSs stem from distinct mutational processes, SBSs are usually caused by lesions or repair mistakes in single-strand DNA, while CNAs are the results of double-strand DNA breaks, double-strand DNA repair defects, DNA replication or cell division defects [3].

CNAs have critical roles in activating oncogenes and in inactivating tumor suppressors [4, 5]. Additionally, aneuploidy status can influence cancer cell proliferation and competitiveness [6, 7]. In morphologically normal tissues, similar SBS have been observed as in cancer cells, however, CNAs are mostly observed in cancer cells but not in morphologically normal tissues [8]. CNAs have been reported to predict cancer relapse and prognosis [9, 10]. These observations suggest that CNAs play a critical role in the malignant transformation of normal cells to cancer cells. However, the underlying mechanism is largely unknown.

Genomic DNA alteration signatures are recurring genomic patterns that are the imprints of mutagenic processes accumulated over the lifetime of cancer cell [11, 12]. Genome alteration signature analysis can not only provide the mutational process information, but also biomarkers for cancer precision medicine [13, 14]. SBS signature analysis has been extensively studied, and represents a prototype for other types of signature study [12]. Despite the importance of CNA in cancer progression, a comprehensive understanding of the mutational process and signature of CNA is still lacking.

Signatures of structural variations (SV) have been studied in breast cancer [15]. However this method relies on high coverage WGS data, and cannot be applied with SNP array or WES data. Macintyre et al employed a mixture modeling based method for copy number component extraction [16]. For each different dataset or cancer type, a different set of CNA components will be generated based on the distributions of the six CNA features (segment size, change point copy number and segment copy number, breakpoint count per 10 Mb, length of segments with oscillating copy number and breakpoint count per chromosome arm). This inconsistency of the CNA components among different datasets limits the generalization of their method in different cancer types or different datasets. Steele et al reported a method for CNA signature analysis with 48 features which considered the absolute copy number, size and heterozygosity status of CNA segment, however the background information of the CNA segment has not been incorporated [17]. We recently developed a predefined set of CNA components and corresponding software implementation as a module of Sigminer (https://cran.r-project.org/package=sigminer) for CNA signature analysis [18-20]. This set of CNA components incorporate the six reported CNA features and include two additional features. The application of this tool (Sigminer) in prostate cancer reveals distinct CNA mutational processes and clinical outcomes [18]. The Macintyre et al method, Steele et al method and our recent method have a limited number of CNA features, a unified and comprehensive CNA classification method across different cancer types is still lacking.

Here we developed a mechanism-agnostic method for copy number segment categorization and signature extraction. Our method incorporates the following information for each DNA segment: absolute copy number, copy number context, segment length, and loss of heterozygosity (LOH) status. The selection of these copy number features was inspired by known patterns of CNA, such as chromothripsis, and whole genome duplication (WGD) [3, 21, 22]. With this new copy number signature analysis method, a pan-cancer landscape of CNA signature is shown. Known CNA patterns have been reproduced, and new CNA signatures have been identified in this pan-cancer study, such as haploid chromosome and combination with WGD. Underlying mutational processes for the identified CNA signatures have been investigated. The activities of some CNA signatures consistently predict cancer patients’ prognosis, suggesting CNA signatures could be cancer prognosis biomarkers.

## Results

### The pan-cancer landscape of copy number alterations

We used the Whole Genome Sequencing (WGS) dataset from Pan-Cancer Analysis of Whole Genomes (PCAWG) to study the profile of CNA. The CNA profiles are then validated with independent The Cancer Genome Atlas (TCGA) SNP array dataset. PCAWG dataset contains the WGS (38-60X sequencing) data of 2778 samples (32 cancer types) [23]. TCGA dataset includes SNP array data (data platform: Affymetrix SNP 6.0) of 10851 samples (33 cancer types) [1].

The length distribution of CNA in the PCAWG dataset and TCGA dataset are shown (Figure 1A). Similar to the previous observation [2], the focal CNAs occur at a frequency inversely related to their lengths, arm-level CNAs occur more frequently than would be expected by the inverse-length distribution associated with focal CNAs. This indicates that compared with focal CNA, chromosome arm-level CNAs are generated through different mutational processes. CNA burden measures the percent of copy number altered genome [9]. Pan-cancer distributions of CNA burden and CNA segment number are shown (Figure 1B and C, Supplementary Figure 1 A and B).

**Figure 1.**
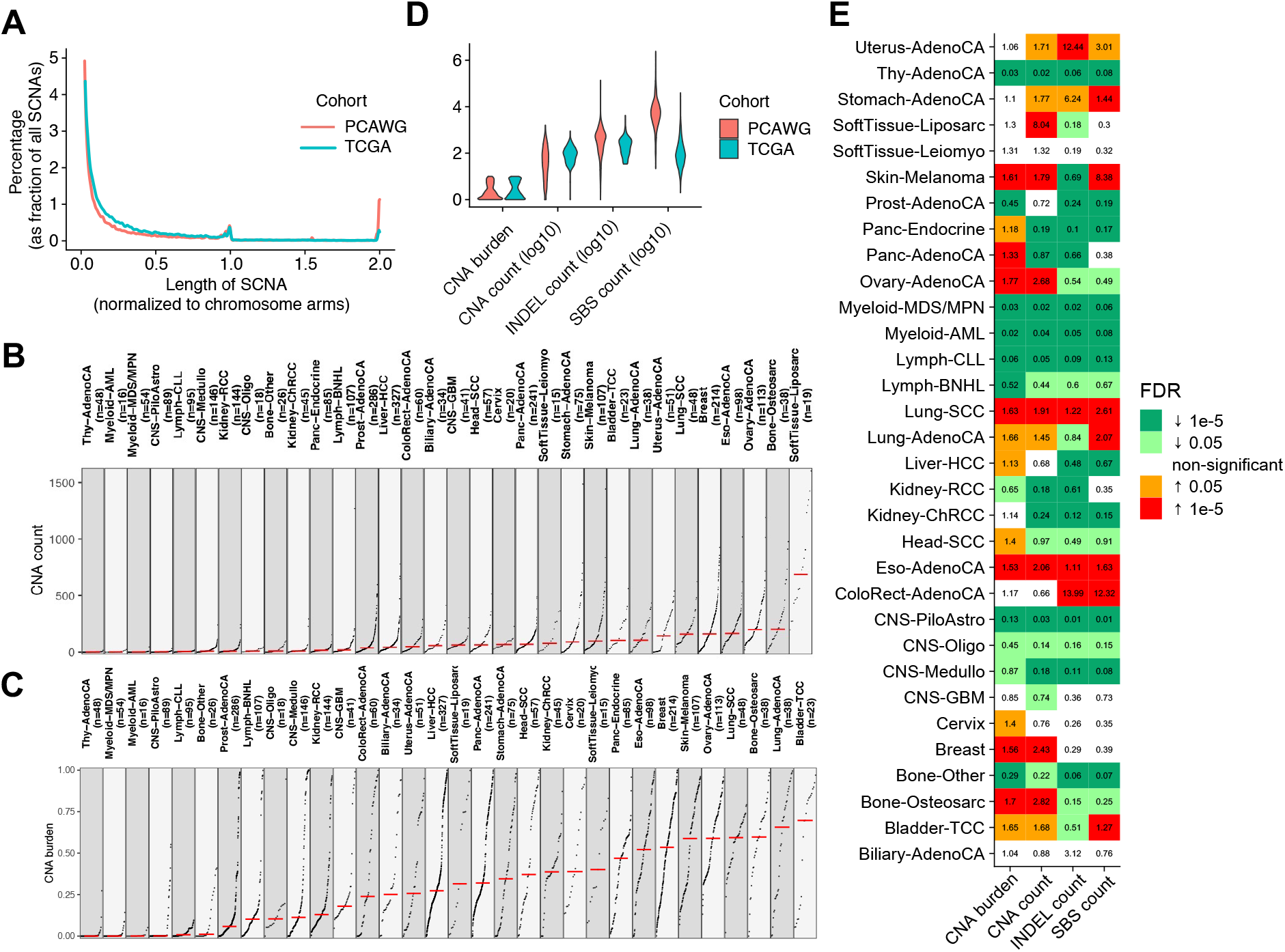
Pan-cancer distribution patterns of CNA. (**A**) Length distribution of CNA in PCAWG (red line) and TCGA (blue line) pan-cancer datasets. (**B, C**) Pan-cancer distribution pattern of CNA count (**B**), CNA burden (**C**) in individual cancer types of PCAWG dataset. (**D**) Pan-cancer distribution of the values of CNA burden, CNA count, INDEL count and SBS count in individual tumors of PCAWG (red) and TCGA (blue) dataset. (**E**) Pan-cancer distribution of the enrichment scores for CNA burden, CNA count, INDEL count and SBS count in PCAWG dataset. The enrichment scores are calculated as the ratio of mean value of specific cancer type compared with the mean value of the whole PCAWG dataset. The colors indicate the FDR corrected *P* values of Mann-Whiney U-test.

Pan-cancer density distribution of the values of CNA burden and CNA count show multi-modal distribution (Figure 1D). In PCAWG dataset, CNA burden and total CNA show a strong positive correlation, SBS and INDEL show a strong positive correlation, CNA and SBS, INDEL show a weak positive correlation (Supplementary Figure 1 C). In TCGA dataset, similar correlations exist as in PCAWG dataset, except, CNA and INDEL show weak negative correlation (Supplementary Figure 1 C). INDELs of TCGA dataset are derived from WES, while INDELS of PCAWG dataset are detected using WGS data, the difference in noncoding regions could contribute to this discordance in CNA and INDEL correlation. Pan-cancer distributions of these genome alteration features (CNA burden, Total CNA, Total INDEL, Total SBS) in PCAWG (Figure 1E) and TCGA (Supplementary Figure 1 D) are shown. Some types of cancer show over-representation of CNA but not SBS, INDEL, such as breast, ovarian cancer, while cancer types including lung cancer show over-representation of all these types of genome alterations (Figure 1E).

### Design of CNA features and components

An essential step for CNA signature analysis is to design proper CNA features and components to classify CNAs. A clearly and stably defined set of CNA components is important for the generalization of CNA signature analysis in various types of cancer. Here we developed a new CNA classification method for CNA signature analysis. Our new classification method considers the following CNA features: morphology or context of CNA, absolute copy number, LOH status. Furthermore, our method is applicable to different types of raw data, including WGS, WES, SNP array or panel sequencing data. Each CNA segment was classified by considering the following detailed features, 1, segment context, including segment shape composed of both the left and right segments of the target segment (low-low, high-high and ladder) and copy number change number. In total, 6 segment context shapes have been defined (Figure 2). 2, Absolute copy number. Including the following components: 0, 1, 2, 3, 4, 5-8 and ≥9. 3, LOH status. 4, segment size, including the following components [24]: S (length<50kb); M (50kb ≤ length<500kb); L (500kb ≤ length<5Mb); E (5Mb≤length). In total, 176 components have been defined to characterize the CNA segments of human cancer patients (Supplementary Table 1).

**Figure 2.**
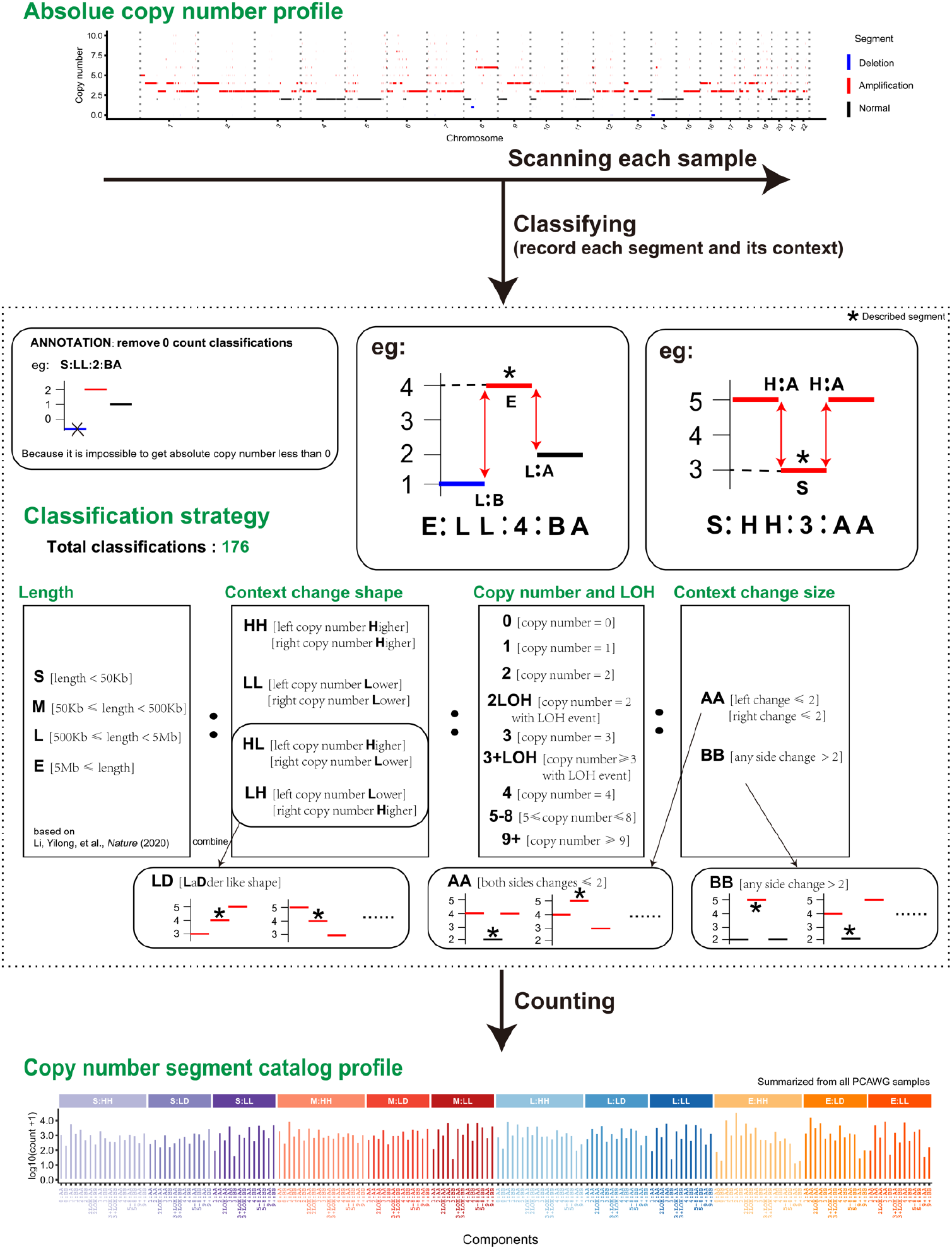
CNA classification strategy for signature analysis. For each CNA segment, the following features have been considered: 1, segment context, including segment shape and copy number change number; 2, Absolute copy number; 3, LOH status; 4, segment size. In total 176 types of CNA segments have been defined accordingly.

### Copy number signature extraction in pan-cancer datasets

Based on the features and components of CNA segment defined above, for each cancer sample, the values for each CNA component will be calculated from the absolute copy number profiles derived from WGS or SNP array data. A copy number component value matrix was generated by combining component values in all tumors. This matrix was subjected to non-negative matrix factorization (NMF), a method previously used for deriving SBS signatures [11]. For de novo copy number signature extraction, we applied the widely used tool Sigprofiler (Supplementary Figure 2) [12]. Sigprofiler has also been used for the extraction of the standard SBS, DBS and ID mutational signatures stored in the COSMIC (catalogue of somatic mutations in cancer) compendium [12].

As reported previously, the choice of the number of mutational signatures is rarely amenable to complete automation [12]. Here, the number of signatures extracted was determined using two parameters. First is the reconstruction error, and the average Frobenius reconstruction error is reported. Second is the stability of signature extraction, and the cosine similarity between the extracted signatures, average silhouette width, is reported (Supplementary Figure 3). In the 2778 PCAWG WGS dataset, 14 CNA signatures have been extracted, these signatures have been named as CNS1, CNS2…CNS14 throughout this study (Supplementary Figure 4). With the building of pan-cancer copy number signature repertoire, for any cancer patient with absolute copy number profile available, we can re-construct the composition of copy number signature through single sample signature fitting (Supplementary Figure 5 and Supplementary Figure 6). Compared with our previous method [18], the new method reported here could reveal potentially unknown patterns, additionally it has the following advantages: 1, the uniqueness of signature profile reflected in the similarity comparison of CNA signature profiles is improved (Figure 3A). 2, the reconstruction error in signature extraction is decreased (Figure 3B). Macintyre et al. applied a mixture modeling based method for copy number component extraction, and the CNA component values are not consistent in different datasets, and this prohibits the signature comparison using cosine similarity analysis across different datasets [16, 18]. These differences between the CNA signatures extracted with Macintyre method, Wang method and this study have been illustrated using PCAWG ovarian cancer dataset (Supplementary Figure 7).

**Figure 3.**
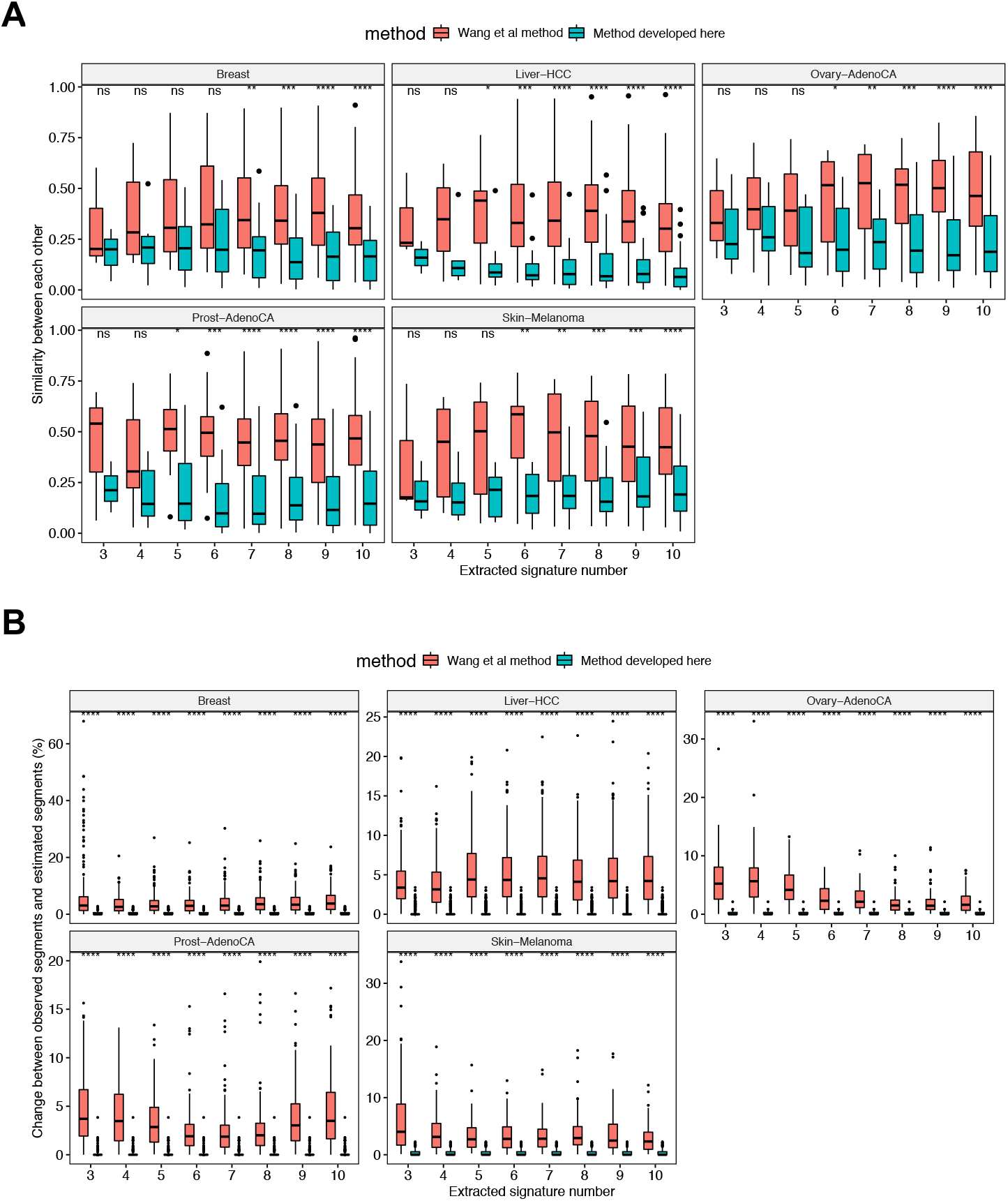
Comparisons between Wang method and the method reported here. (**A**) Inter-signature similarity comparison between Wang method and the method reported here. In five independent PCAWG cancer sub-datasets, consistently lower inter-signature similarities are observed with the method described here. (**B**) Comparison of signature reconstruction error between Wang method and the method reported here. The method reported here show lower signature reconstruction error. Wilcoxon rank-sum test was performed to test for the differences between the two groups. ns: *P* > 0.05, *: *P* ≤ 0.05, **: *P* ≤ 0.01, ***: *P* ≤ 0.001, ****: *P* ≤ 0.0001.

### Benchmark analysis of CNA signatures

To evaluate the robustness of the proposed copy number signature analysis procedure, we compared the CNA signatures extracted in 2778 PCAWG WGS dataset with the CNA signatures derived from 10851 TCGA SNP array dataset (Supplementary Figure 4 and Supplementary Figure 8). In 10851 TCGA SNP array data, 20 CNA signatures have been extracted, these TCGA CNA signatures have been named as Sig1, Sig2…Sig20 throughout this study (Supplementary Figure 8). The similarities between signatures extracted from TCGA SNP array dataset and PCAWG WGS dataset are calculated using cosine similarity analysis method (Figure 4A). Most (9/14) PCAWG signatures can have highly similar (cosine similarity R≥0.8) counterparts in TCGA dataset. Four PCAWG signatures (CNS4, CNS5, CNS6, CNS14) have intermediate similarity (R≥0.51) counterparts in TCGA dataset. Majority of TCGA CNA signature (16/20) have median to high similar counterpart signature in PCAWG CNA signature set. Four TCGA CNA signatures (Sig15, Sig18, Sig9, Sig20) do not have matched PCAWG signature. All these four un-matched TCGA CNA signatures are tissue specific signatures. Using 10% relative activity as a cut off, TCGA Sig20 is only observed in TCGA testicular germ cell tumors (TGCT), and this type of cancer is not included in PCAWG dataset. TCGA Sig18 is only observed in TCGA thymoma (THYM) cancer type, which is also not included in PCAWG dataset. TCGA Sig15 is majorly observed in adrenocortical carcinoma (ACC), and to a less extent in kidney chromophobe (KICH) carcinoma, and ACC is not included in PCAWG dataset. TCGA Sig 9 is observed in ACC and TGCT, both ACC and TGCT are not included in PCAWG dataset.

**Figure 4.**
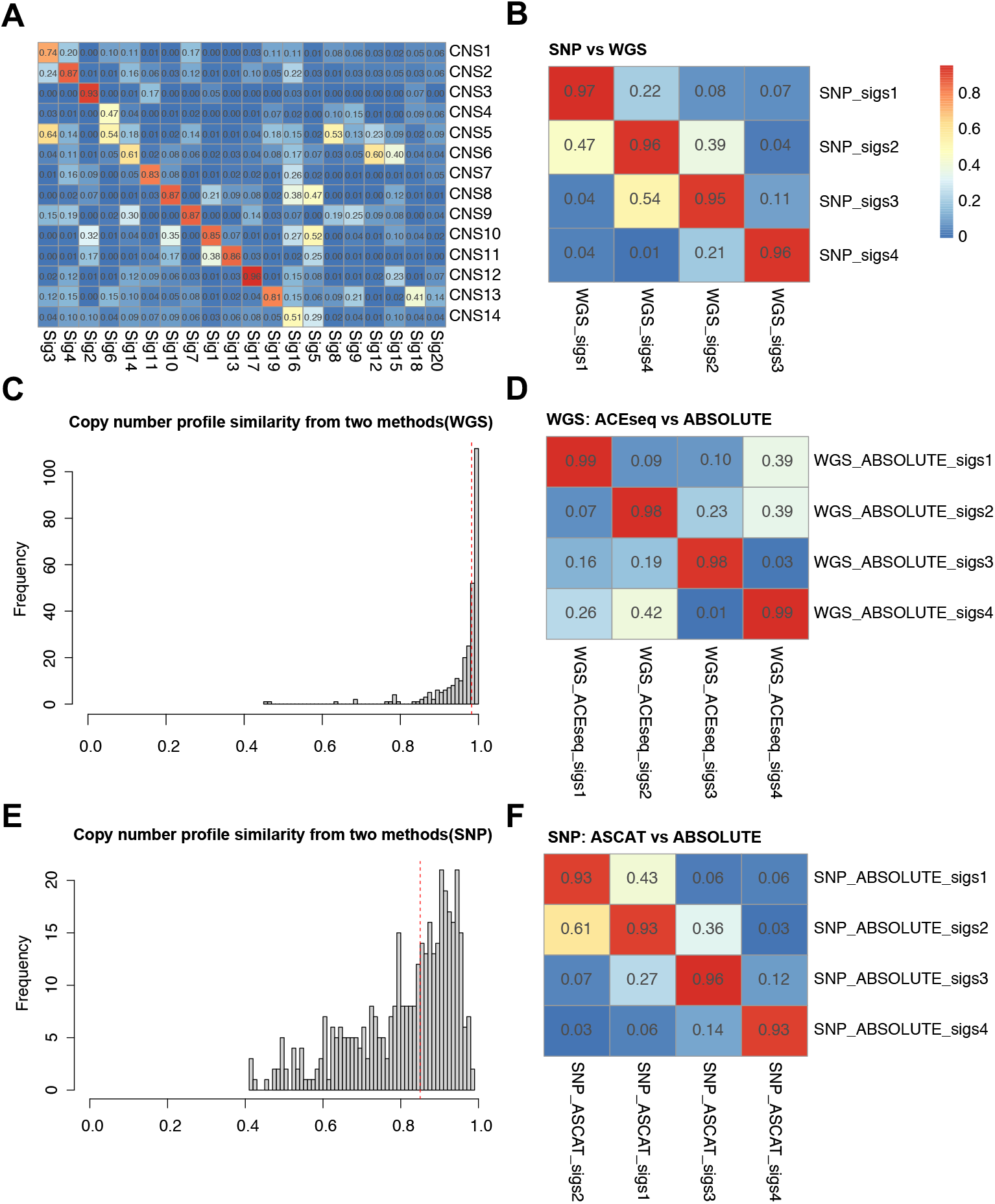
CNA signature benchmark analysis with CNA profiles derived from different platforms and different CNA calling algorithms. (**A**) Inter-correlations between the CNA signatures extracted in PCAWG dataset and TCGA dataset. Cosine similarity values are reported for each comparisons. (**B**) Cosine similarities are reported in comparing CNA signatures extracted from WGS platform and SNP array platform. (**C**,**D**) CNA profiles have been extracted from 286 WGS samples using ACEseq or ABSOLUTE algorithm, distribution of copy number profile cosine similarities between the two methods is reported (**C**), pairwise comparisons of the CNA signatures extracted with the CNA profiles called with the two algorithms are shown (**D**). (**E**,**F**) CNA profiles have been extracted from 468 SNP array samples using ASCAT or ABSOLUTE algorithm, distribution of copy number profile cosine similarities between the two methods is reported (**E**), pairwise comparisons of the CNA signatures extracted with the CNA profiles called with the two algorithms are shown (**F**).

The fact that CNA signatures extracted de novo from PCAWG (WGS based) and TCGA (SNP array based) show a pairwise similar signature profile suggests the robustness of our method in identifying true cancer patterns. To further evaluate the performance of our method, we carried out benchmark analysis with CNA profiles derived from different platforms and different CNA calling algorithms. For benchmark analysis with CNA data from different platforms, CNA signatures have been extracted independently from 286 WGS and 468 SNP array derived prostate cancer CNA profiles. The CNA signatures extracted from WGS data are highly similar to the CNA signatures extracted from SNP array data (median cosine similarity R=0.96) (Figure 4B). The effects of different CNA calling algorithms on the stability of CNA signatures have been evaluated (Figure 4C and F). With WGS, the signatures derived from CNA profiles called with ABSOLUTE [25] algorithm are highly similar to the signatures derived from the CNA profile called with ACEseq algorithm (median cosine similarity R=0.98) (Figure 4C and D). Similarly, for SNP array platform, the CNA signatures derived from ABSOLUTE algorithm are highly similar to the CNA profile derived from ASCAT [26] algorithm (median cosine similarity R=0.93) (Figure 4E and F). In conclusion, the CNA signature extraction method proposed in this study can be applied to CNA profiles derived from different platforms and different CNA calling algorithms, and the CNA signatures extracted with our algorithm are stable, and could reflect the true DNA alteration patterns of cancer.

### Pan-cancer distribution of CNA signature

The proportion of tumors with the signature and median activity of the signature in different types of cancers are shown for PCAWG dataset (Figure 5A). CNS3 was observed in tumors with few CNA count, such as AML (Figure 5A and Supplementary Figure 9). The enrichment scores (defined as the ratio comparing the mean value of specific cancer type vs mean value of pan-cancer dataset) of the activities of CNA signatures in different types of cancer are shown for PCAWG dataset (Figure 5B). Some CNA signatures show enrichment in specific cancer type, for example, the enrichment score of CNS4 in PCAWG liposarcoma is 52.91 (Figure 5B and Supplementary Figure 10). The profile of CNS4 suggests the presence of extrachromosomal DNA (ecDNA) or neochromosome. Actually, double minutes, small, self-replicating extrachromosomal structures in a ring form, were originally observed in sarcomas [27]. The presence and evolution of neochromosome has also been investigated in liposarcomas [28]. Pan-cancer distributions of the relative and absolute activities of CNA signatures are shown for PCAWG dataset (Figure 5C and D). Compared to other signatures, CNS3 has the highest relative activities. Pan-cancer profiles of TCGA CNA signatures are shown (Supplementary Figure 11).

**Figure 5.**
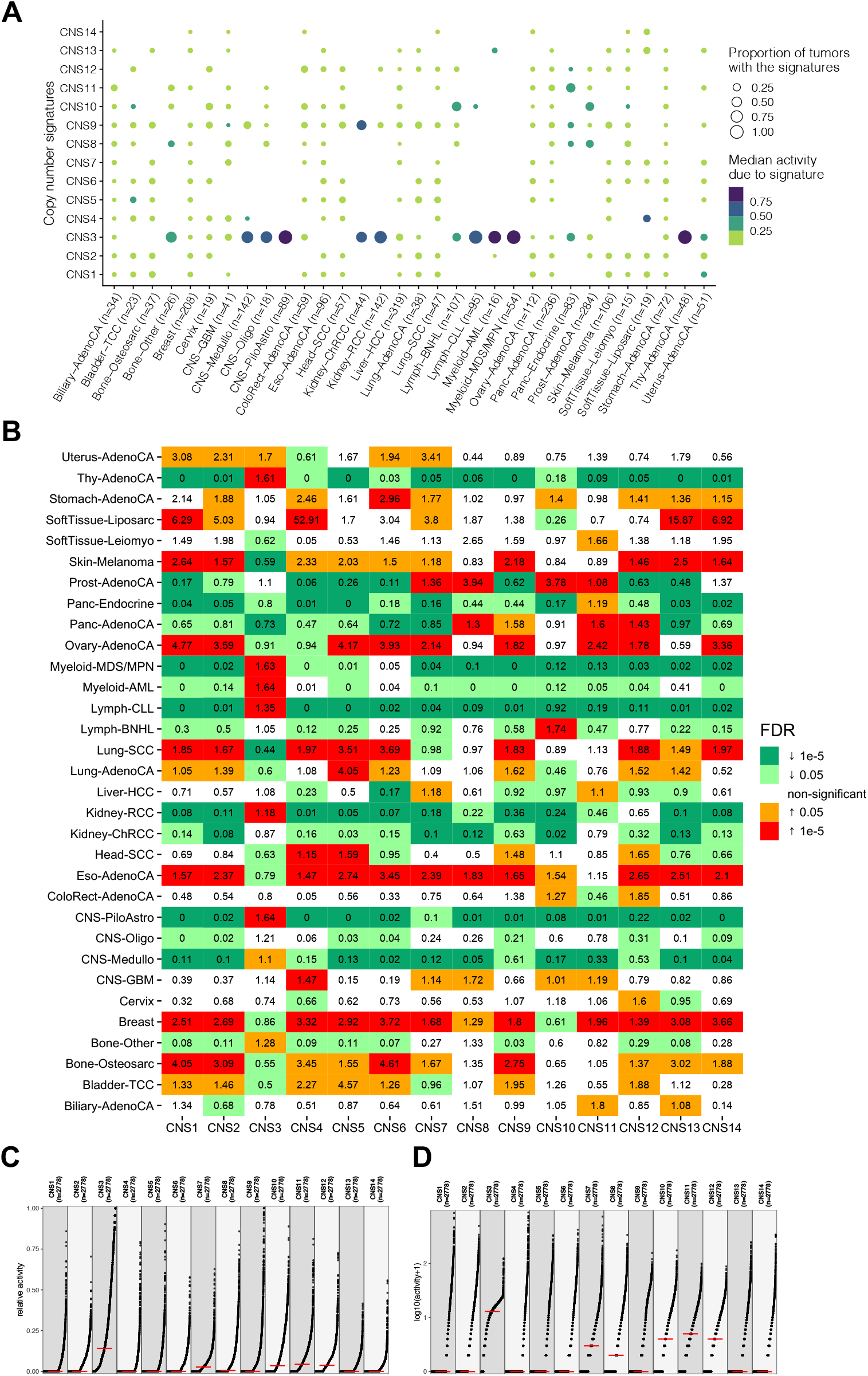
The activity distribution of CNA signatures in PCAWG pan-cancer dataset. (**A**) Proportion of tumors with the signature and the median activity of the signature are shown for 32 PCAWG cancer types. For each individual tumor, only signatures that contribute to ≥5% of the total are shown. (**B**) Enrichment score analysis of CNA signature in PCAWG dataset. The enrichment score are calculated as the ratio of mean signature activity of specific cancer type compared with the mean signature activity of the whole PCAWG dataset. The colors indicate the FDR corrected *P* values of Mann-Whiney U-test. (**C**,**D**) Relative activity (percentage of the total) (**C**) and absolute activity (contributed CNA segment number) (**D**) distribution pattern of CNA signatures in PCAWG dataset are shown.

Compared with SBS, DBS, ID signature profiles, the profile of CNA signature show much reduced inter-signature correlation, suggesting an increased signature distinctness compared with other types of signature profiles (Figure 6A and B). In PCAWG dataset, the average number of CNA signatures in a single patient is approximately 2 (Figure 6C). Some cancer types have few CNA signature, such as AML. Some have many CNA signatures, such as breast cancer and ovary cancer (Figure 6D). This is in line with the fact that AML has a low number of CNA segment counts, while breast cancer and ovarian cancer have many CNA segment count (Figure 1B). Several known copy number patterns can be reproduced in our study using PCAWG dataset. For example: stable genome (CN-Sig 5 in Wang et al 2021 study) [18]; ecDNA; chromothripisis; WGD, homologous recombination deficiency (HRD). Several new patterns have been identified in this study, including the following (Supplementary Figure 4 and Supplementary Figure 12): Pattern 1: Focal homozygous deletion (CNS10). This pattern is featured with regions of homozygous deletion. Generally homozygous deletion is focal, and surrounding tumor suppressor genes. Pattern 2: Haploid chromosomes (CNS11). This pattern is characterized by a mixture of chromosome with copy number 1 and 2. This haploid status of several chromosomes could be caused by cell cycle defects. In the published literature, we can find that some cancer cells are haploid, and this could be derived through similar mechanism [29, 30]. Pattern 3: Haploid chromosome and WGD (CNS6). This pattern is featured with chromosome LOH with copy number 2 or 3. This pattern could be formed through haploid chromosome then WGD.

**Figure 6.**
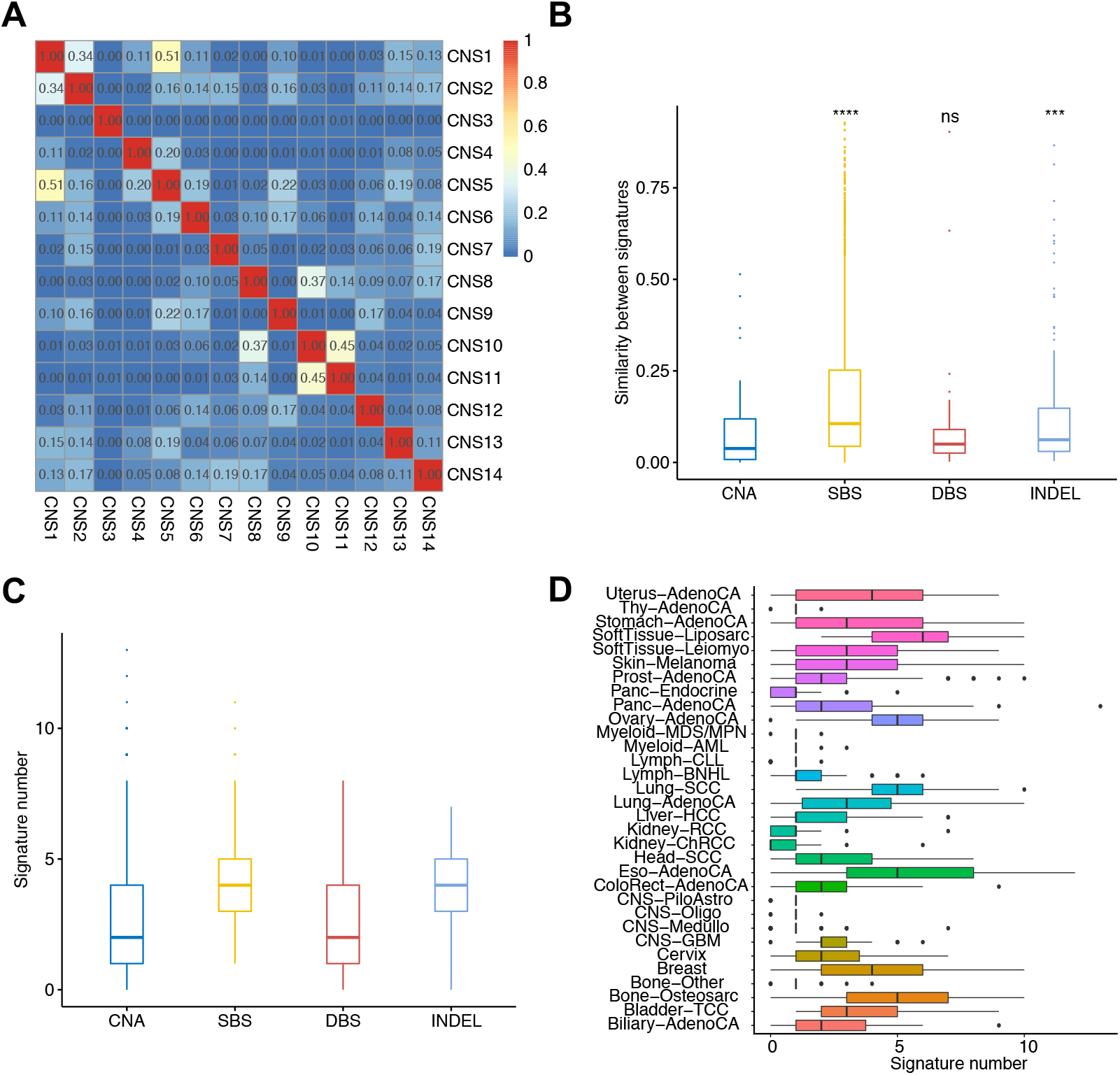
Distinctness and the number of CNA signatures extracted in PCAWG dataset. (**A**) Inter-correlation analysis of the profiles of 14 PCAWG CNA signatures. The numbers are cosine similarity values comparing each pairs of CNA signatures. (**B**) Inter-signature similarities comparing the signatures of CNA, COSMIC reference SBS, DBS and ID (insertion and deletion) signatures. (**C**) Median number of signatures for CNA, SBS, DBS, INDEL in PCAWG dataset. (**D**) Distribution of the number of CNA signature in each cancer types of PCAWG dataset.

### Potential mutational processes for CNA signatures

An important purpose of CNA signature analysis is to identify the underlying mutational processes for CNA. Potential mutational processes for CNA include intrinsic inducers and extrinsic inducers. Intrinsic CNA inducers include: double-strand break repair defects (HRD etc.); cell cycle defects; DNA replication defects; telomere loss, etc. Extrinsic CNA inducers include chemical or physical agents that induce double-strand breaks, or interfere with the cell cycle, such as chemotherapy drugs or ionizing radiation. Smoking and ultraviolet (UV) are known to induce SBS signatures, however no specific CNA signatures are associated with smoking and UV (Supplementary Figure 13 and Supplementary Figure 14), probably because smoking and UV induce single-strand lesions, and are not associated with double-strand break, and the consequent CNA.

We group annotations of pathogenic germline variants and somatic driver mutations in DNA-repair genes across the PCAWG dataset, then correlate their presence with activities of the copy number signatures (Figure 7A). *BRCA1* functional mutations are significantly enriched in CNS14, suggesting the presence of HRD. CNS3 is significantly associated with patients without *TP53* mutation, suggesting a state of stable genome. COSMIC SBS3 and ID6 are known HRD associated signatures, they also show strong correlations with CNS14 (Figure 7B). Associations between the presences of cancer driver mutations with the activities of copy number signatures are shown for PCAWG and TCGA dataset (Supplementary Figure 15 and Supplementary Figure 16). Some driver mutations are significantly enriched in specific CNA signature. For example *SPOP* mutation is specifically associated with CNS8 in prostate cancer (Supplementary Figure 15 and Supplementary Figure 17).

**Figure 7.**
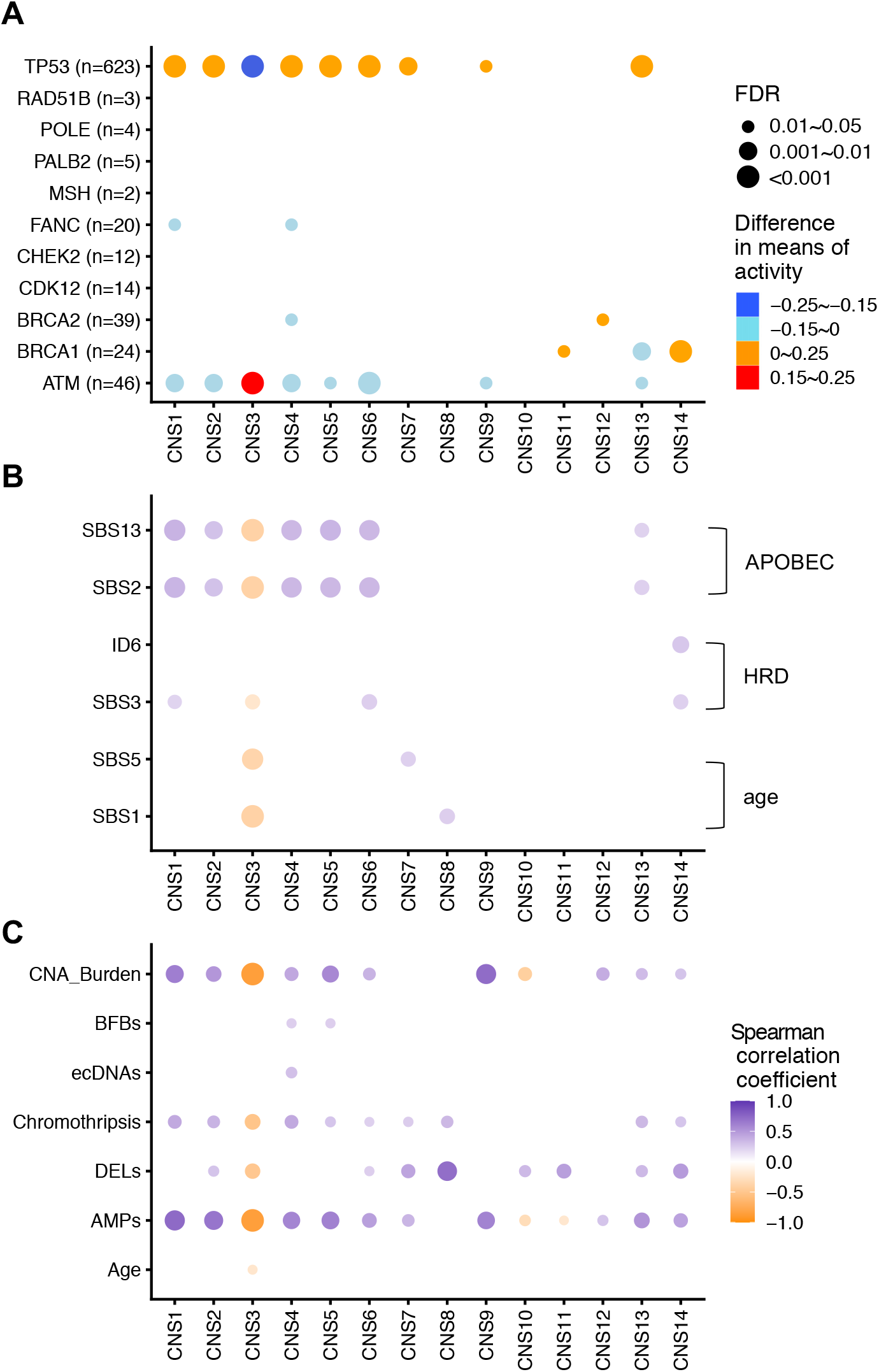
Correlations between CNA signatures and different types of genome alterations. (**A**) Associations between gene mutation status in key DNA repair genes with the relative activities of CNA signatures. Association of pathogenic mutations (germline and somatic combined) in key DNA repair genes with the relative activities of CNA signatures. For each gene, PCAWG patients are divided into two groups based on the mutation status of the gene. The difference values in mean relative activities between two groups (activities of mutated group – activities of un-mutated group) are reported, and FDR corrected *P* values (Wilcoxon rank-sum test) are shown. The color and size of the points represent the differences and adjusted *P* values, respectively. MSH refers to MSH2, MSH3, MSH4 and MSH6, genes in the mismatch repair pathway; FANC refers to genes associated with Fanconi anaemia, namely FANCA, FANCC, FANCD2, FANCE, FANCF, FANCG, FANCI, FANCL and FANCM. The number after gene or gene cluster name refer to the number of pathogenic mutations (germline and somatic combined). (**B**) Inter-correlations between the activities of CNA signature and APOBEC, homologous recombination deficiency (HRD), age signatures in PCAWG dataset. (**C**) Associations between the relative activities of each signature and the indicated cancer genome features are calculated from individual cancer patient. Spearman correlation coefficient values are reported.

Correlations between some cancer genome features, such as CNA burden and the presence of ecDNA, with the activities of CNA signatures are displayed for PCAWG dataset (Figure 7C). In association analysis, activity of CNS9 is the most accurate predictor for WGD, suggesting CNS9 could be a reflection of the status of WGD. CNS14 activity is the most accurate predictor for HRD, suggesting CNS14 could be the major signature of HRD (Supplementary Figure 18). The activities of most CNA signatures show significant tumor stage difference, for example, CNS4 is present in higher level in stage III than in stage I tumors (Supplementary Figure 19). Representative sample profile, notable features and potential mechanisms or mutational processes for PCAWG and TCGA CNA signatures are summarized respectively. (Supplementary Figure 12 and Supplementary Figure 20). The mechanisms for several CNA signatures are still unknown.

### The clinical relevance of CNA signatures

The CNA signatures extracted from cancer patients could be cancer prognosis biomarkers. To test this hypothesis, Cox regression analyses were conducted to evaluate the associations between the activity of each CNA signature and cancer patients’ overall survival time for each cancer types. For each Cox model, we report a Z-score that encodes the directionality and significance of the survival relationship. A Z-score of >1.96 indicates that the upregulation of the target feature (activity of CNA signature) is related to the reduction of the survival time at the *P*<0.05 threshold, while the Z-score of <-1.96 indicates that the increase in the target feature at the *P*<0.05 threshold will indicate a longer survival time.

To compare the prognostic effects of CNA signature activity in TCGA and PCAWG dataset, we select the cancer types that have sufficient number of patients (n>50) with both CNA signature activity and OS data available in both TCGA and PCAWG dataset. We calculate the CNA signature activity of both TCGA and PCAWG dataset using single sample fitting with TCGA signature set (Sig1, Sig2…Sig20). In total kidney cancer, stomach cancer, liver cancer, ovarian cancer and melanoma sub-datasets are available for this prognosis comparison analysis. Z-scores of CNA signatures are compared in pair in matching TCGA and PCAWG cancer types (Figure 8A). The Z-scores of CNA signature activity in matching TCGA and PCAWG cancer types are significantly correlated (Spearman correlation R=0.44, *P*=0.015), suggesting the robustness of CNA signature in predicting cancer patients’ overall survival (Figure 8B). For example, in kidney cancer, high CNA Sig1 activity is associated with significantly poor overall survival in both TCGA and PCAWG datasets (Figure 8C).

**Figure 8.**
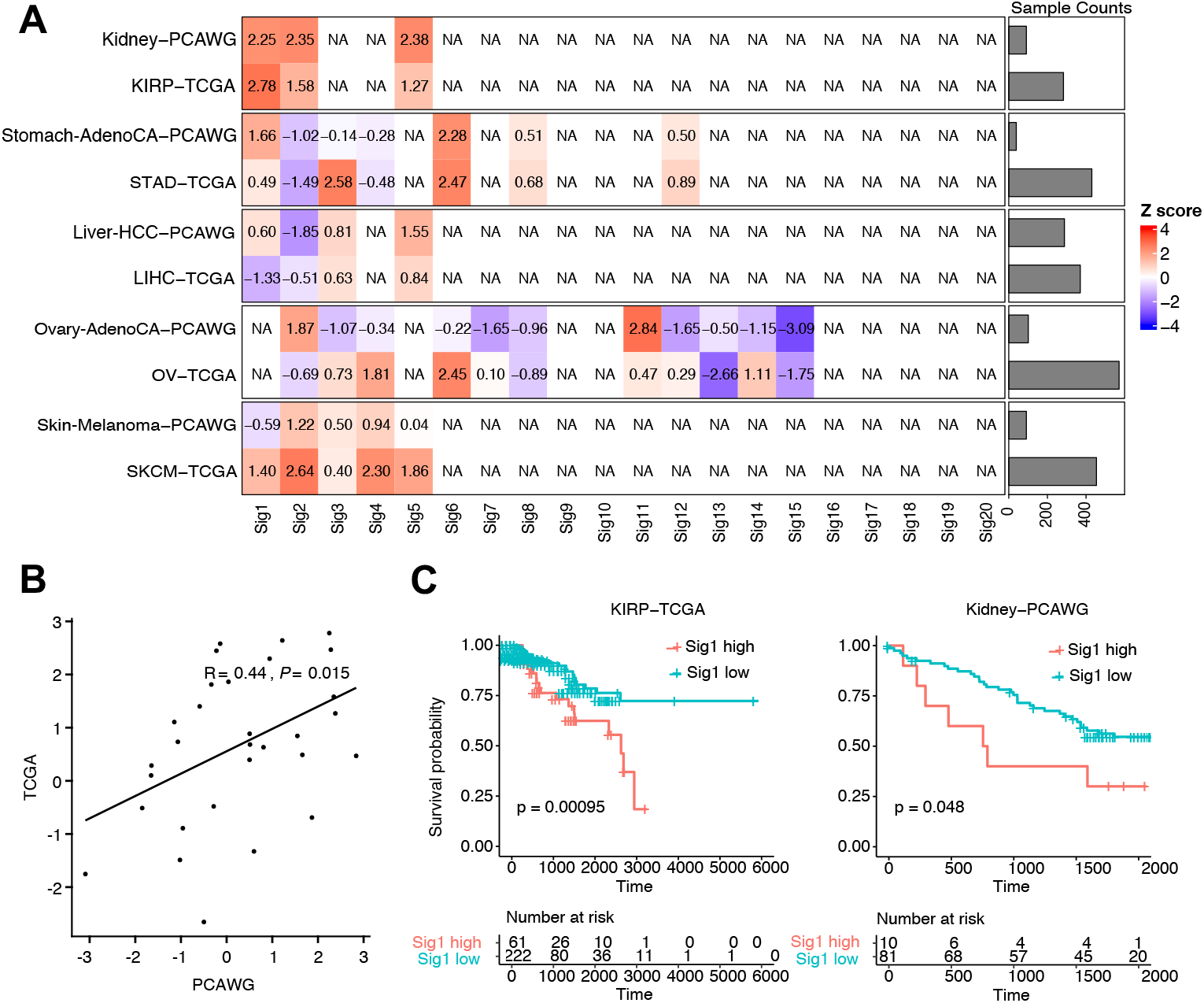
CNA signature activity and cancer patients’ prognosis. (**A**) The prognosis Z-scores of the activity of CNA signatures are calculated in each matching TCGA or PCAWG cancer types with sufficient number of patients (n>50) having overall survival and CNA profiles available for this analysis. NA indicates the CNA signature does not exist in the specific cancer type; colors reflects the values of Z-scores. (**B**) Correlations between the Z-scores of TCGA and PCAWG matching cancer types. Pearson test *P* value is reported. (**C**) The activity of Sig1 shows consistent prognosis in the kidney cancer of TCGA (left) and PCAWG (right) dataset. Kaplan-Meier overall survival curves show the comparison between different groups stratified by CNA signature Sig1 activity. Samples with Sig1 activity higher than the cutoff (determined by surv_cutpoint function of “survminer” package) were classified as “Sig1 high” group, and samples with Sig1 activity less than the cutoff were classified as “Sig1 low” group. The log-rank test *P* values are reported. KIRP: Kidney renal papillary cell carcinoma; STAD: Stomach adenocarcinoma; LIHC: Liver hepatocellular carcinoma; OV: Ovarian serous cystadenocarcinoma; SKCM: Skin Cutaneous Melanoma.

## Discussion

Here we developed a unified and comprehensive method for copy number signature analysis. Our method can be applied in cancer patients with copy number profiles generated with WGS or SNP array data. Our copy number signature analysis method is based on a novel and comprehensive method to catalog copy number segments. New CNA patterns have been identified, and the activities of some CNA signature have been demonstrated to be associated with cancer patients’ prognosis. It will be of interest to investigate the potential application of CNA signatures in predicting the clinical response of certain cancer treatments for example PARP inhibitor treatment or cancer immunotherapy.

Mutational signature analysis is initially developed with SBS, and CNA is different from SBS in several aspects. SBS and CNA are derived from different mutational processes, some common SBS inducers such as smoking and UV do not induce specific CNA signature. Smoking and UV can generate single strand DNA lesions, while CNA is the consequence of DSB or cell division defects [3]. The scale of the influenced genome is different between CNA and SBS. CNA could affect a much larger portion of the genome than SBS. Each SBS is usually a consequence of a unique mutational process, while each CNA segment could be the consequence of multiple CNA mutational processes. Some CNA inducers have impact on global CNA pattern, such as inducers of cell division defects leading to WGD or aneuploidy, which have a global impact on CNA profile. The resulting signatures of global CNA mutational processes need to be combined with other CNA mutational processes, leading to new signatures. For example, WGD and HRD combination generate new signature, which cannot be reconstructed as a linear combination of single WGD and single HRD signature. This suggested that the mutational processes for the CNA signatures reported in this study can be combinations of different processes. For example, CNS6 could be a combination of haploid chromosome (CNS11) and WGD (CNS9). The evolution timeline for mutational process of CNA is largely unknown. With the availability of cancer cell fraction (CCF) information for CNA segments [25, 31], we can calculate the activities of CNA signatures for clonal CNA and subclonal CNA, and this analysis can provide distinct insight into the evolution of CNA mutational processes and CNA patterns (Supplementary Figure 21).

Global patterns and mutational processes for CNA in human cancer are largely unknown. Currently known methods for CNA signature analysis include Macintyre method and Wang method, and both methods classify CNA segments using known patterns, such as the lengths of oscillating copy number segment chains (named “OsCN”) [16, 18]. Some unknown CNA patterns that do not fit into these known patterns cannot be detected with these approaches. Here we provide a mechanism-agnostic method for CNA signature analysis, this method can reveal potentially new CNA patterns, and is suitable for pan-cancer study. For copy number signature analysis, detailed sequencing information is not required. Compared with previously reported SV signature [15], the advantage of this method is the wider application area, for example, panel sequencing data or SNP array data could be used to derive CNA signature. The disadvantage is the increased uncertainty. Sometimes two nonequivalent genomes can produce exactly the same copy number patterns. The underlying mutational process for copy number pattern/signature could be complicated by these uncertainties.

During the submission process of this manuscript, Steele et al reported pan-cancer copy number alteration signature using a different mechanism-agnostic approach [17]. Compared to Steele et al method, this study incorporates the context or shape information of each CNA segment, and this can reveal additional insight into the mutational processes and patterns of CNA. These CNA shape information include “ Low-Low “ “High-High”, “Ladder” and also the extent of copy number change (>2 or ≤2) (Figure 2). Thus our methods can reveal additional insights into CNA patterns and mutational processes compared with Steele et al method. Some patterns of copy number alteration can only be detected with our method but not Steele et al method. For example the oscillation status of CNA segment, the extent of copy number change flanking specific CNA segment. The incorporation of this additional CNA segment shape or context information enable us to directly interpret the mechanism of CNA signature, while Steele et al study totally rely on association study for CNA signature mechanism interpretation. Furthermore our CNA signature profiles have been constructed and compared in two different pan-cancer datasets: TCGA SNP array dataset and PCAWG WGS dataset. The prognostic effects of CNA signatures have also been compared and validated in the above two different datasets, while Steele et al only performed analysis in TCGA dataset. Compared to recent publication, our study provide an innovative and enhanced approach for CNA segment categorization and signature analysis, and reveal distinct insight into the patterns, evolution, mutation process of CNA.

CNA signatures have just started to be investigated. Many issues still remain to be studied. One of the major purpose of CNA signature analysis is to reveal the underlying mutational process for CNA. The mutational processes for several CNA signatures reported in this study are unclear. This needs further association studies, and also experimental studies with pre-defined CNA inducers. Due to the heterogeneity of tumor, the final CNA profile can be a combined results of many heterozygous CNA difference. Single cell CNA signature analysis could reveal the heterogeneity and evolution process of cancer.

### Conclusions

Global patterns and mutational processes for CNA in human cancer are largely unknown. Here we developed a method to reveal pan-cancer patterns of DNA copy number alteration signatures through a mechanism-agnostic approach. Our results correlate some CNA signatures to known biological characteristics through diverse approaches ranging from signature profile observation to molecular profiling. New CNA patterns have been identified, and the activities of some CNA signature have been demonstrated to be associated with cancer patients’ prognosis. Collectively, this method paves the way for further study on revealing new CNA etiology and designing robust biomarkers for cancer precision diagnosis and therapy.

## Materials and Methods

### Copy number calling and processing

PCAWG allele-specific copy number generated by PCAWG-11 working group are the result of a bespoke procedure that combines output from 6 different copy number callers: ABSOLUTE, ACEseq, Battenberg, CloneHD, JaBbA and Sclust. First ran all methods across all samples with the consensus SVs included and applied an algorithm across the segmentations to obtain consensus breakpoints. With these mandatory breakpoints the methods were rerun without calling any additional breakpoints. TCGA allele-specific copy number data were generated from Affymetrix Genome-Wide Human SNP 6.0 (SNP6) array with ASCAT2 workflow. The PCAWG and TCGA allele-specific copy number (CN) data values are integers. Copy number alterations were generally classified as such:

- Amplification: segment total CN>2.
- Homozygous deletion: segment total CN=0.
- Loss of heterozygosity: segment with minor CN=0 and total CN>0 and size>10Kb.
- Deletion: segment total CN<2.

### Copy number segment classification

Copy number segments were classified as normal, amplification or deletion. They were further classified by size of the segment (small (S), <50Kb; middle (M), 50Kb-500Kb; large (L), 500Kb-5Mb; extreme large (E), >5Mb). They were then further classified by the context of a segment (the shape), i.e. how total copy number of segment on its left/right side compared to it (HH, left copy number Higher & right copy number Higher; LL, left copy number Lower & right copy number Lower; HL, left copy number Higher & right copy number Lower; LH, left copy number Lower & right copy number Higher). LH and HL have the same shape, so they are combined into LD (Ladder like shape). The segments were further classified by checking if they harbor LOH or not, and their absolute copy number. For LOH segments, the total copy numbers were classified into 1, 2LOH, 3+LOH; for non-LOH segments, the total copy numbers were classified into 0, 2, 3, 4, 5, 6, 7, 8, 9+. Finally, a total of 176 mutually exclusive categories (described as components in this study) were cataloged. Copy number profile of each sample can be inputted into the classification algorithm above and each component will be counted to generate an integer vector. For multiple samples, a component-by-sample matrix will be generated and used for copy number signature discovery. The data import and classification procedure were implemented as functions “readcopynumber” and “sigtally” in R package Sigminer (https://cran.r-project.org/package=sigminer).

### Copy number signature extraction

Copy number signatures were extracted from component-by-sample matrix with golden standard tool SigProfiler v1.0.17 (https://github.com/AlexandrovLab/SigProfilerExtractor) with default parameters. Briefly, this de novo signature extraction includes the following six steps: dimension reduction, resampling, NMF, iteration, clustering and evaluation [11]. For each NMF, the initialization was with random numbers, and iterations were performed for 10,000 – 1,000,000 times until stable results are obtained. This NMF process was repeated for 100 times with resampling data. Clustering the decomposition matrixes to identify the number of signatures from 2 to 30. The SigProfiler has been successfully applied to TCGA and PCAWG pan-cancer data for multiple mutation types including SBS, DBS and INDEL. Two key parameters for determining signature number, stability measured by average silhouette, and the average Frobenius reconstruction error were obtained from the result of SigProfiler. We selected the signature extraction solution as the maximum signature number which meets the following criteria:

1. No over fit.
2. Stability should be at least local maximal.
3. Mean cosine distance should be as small as possible.

Based on the rules, we selected 14 signatures for PCAWG copy number data and 20 signatures for TCGA copy number data. The profile and activity for each signature were obtained, accordingly.

### CNA signature benchmark analysis

We select data from prostate cancer to evaluate the reproducibility of our method with benchmark analysis. 468 SNP-array data is derived from TCGA, we used the ASCAT [26] and the ABSOLUTE [25] algorithm to generate allele-specific copy number. 286 whole-genome sequencing (WGS) data is derived from PCAWG, and ACEseq [32] and the ABSOLUTE [25] algorithm were used to obtain absolute copy number profiles. Copy number segments were categorized into 176 classes as described above in each sample. The cosine similarities between CNA profiles are calculated according to this 176-element vector. First, we benchmarked the impact of platforms on CNA signature analysis. Four CNA signatures are independently extracted from WGS, WES, or SNP array derived CNA datasets with our method described above. Pairwise comparisons between the four CNA signatures are reported. Next, we benchmarked the impact of different CNA calling algorithms in CNA signature analysis. ACEseq and the ABSOLUTE algorithm were applied to obtain absolute copy number profile from 286 WGS data independently, and four CNA signatures have been independently extracted, compared and cosine similarity values of pairwise comparisons are reported. Similarly ASCAT and the ABSOLUTE algorithm were applied to obtain absolute copy number profile from 468 SNP-array dataset independently, and four CNA signatures have been independently extracted and compared.

### Copy number signature labelling and matching

We sorted all copy number signatures based on their total activities to all samples. PCAWG data analysis is the major focus of this study. For PCAWG, we named 14 copy number signatures from CNS1 to CNS14. For TCGA, we firstly named 20 copy number signatures from Sig1 to Sig20. We further added extra labels to their names for better comparing the copy number signatures between PCAWG and TCGA by following the rules:

1. We classified signature similarity based on cosine similarity values into four levels: ≥ 0.8 (High, H), ≥ 0.51 & <0.8 (Intermediate, M), and <0.51 (unmatched).
2. The results were combined for a TCGA signature matched to 2 or more PCAWG signatures. i.e., TCGA Sig3 were matched to PCWG CNS1 and CNS5 both in middle level similarity, so the signature was labelled as “Sig3-CNS1(M)/CNS5(M)”.
3. A postfix with the matched similarity rank was used if a PCWAG signature was matched to 2 or more TCGA signatures. i.e. TCGA Sig8 was labelled as “Sig8-CNS5(M)_3”, here 3 means this signature was the 3rd signature matched to CNS5.

### Group enrichment analysis

To comprehensively show the enrichment of a variable across cancer types, inspired by enrichment analysis in Maftools, we designed and implemented the group enrichment analysis as functions “groupenrichment” and “showgroup_enrichment” of R package Sigminer. To illustrate how this analysis works, here we use CNA burden analysis for PCAWG breast cancer as an example. Firstly, we divided all PCAWG samples into two categories: Breast and non-Breast. Then we compared the means of CNA burden with Wilcoxon rank-sum test and calculated the ratio of the means, i.e., 1.56 means the average CNA burden in Breast cancers is 56% higher in non-Breast cancers. The result of statistical test indicates if this result was randomly obtained. We used heatmap to visualize the final results and distinguished the different results based on both the mean ratio and statistical test result:

- If the comparison result is statistically non-significant, white color was used to fill the heatmap cell.
- If the ratio is bigger than 1 and the comparison result is statistically significant, red color was used to fill the heatmap cell.
- If the ratio is smaller than 1 and the comparison result is statistically significant, green color was used to fill the heatmap cell.

### Association analysis

Associations between the activities of signatures, associations between signature activity and gene mutation status were performed using one of two procedures: 1) for a continuous association variable, Spearman correlation was performed; 2) for a binary variable, patients were divided into two groups and a Wilcoxon rank-sum test was performed to test for differences in average activities of signatures between the two groups. Associations between each CNA signature and WGD or HRD status were performed using a two-sided Fisher’s exact test. All reported p-values were FDR corrected. Only associations with both P ≤ 0.05 and odds ratio > 1 were reported. Full correlation network for continuous variables was constructed using R package “correlation” (https://cran.r-project.org/package=correlation).

### Survival analysis

Associations between copy number signature activities with overall survival were identified using univariate Cox proportional hazard models in each cancer types. For each Cox model, a Z-score that encodes the directionality and significance of the survival relationship is reported. Z-scores reflect the normalized deviations from the mean of a normal distribution, and these Z-scores are calculated following similar procedures as previously described [33]. Briefly, the Cox model is given by:

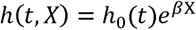

Where *t* is the overall survival time, *h(t,X)* is the hazard function, *h*_*0*_*(t)* is the baseline hazard. *X* is a potential prognostic variable. Z-score is calculated by dividing the regression coefficient β by its standard error.

The prognosis effects of the activity of CNA signatures are compared in matching TCGA and PCAWG cancer types with sufficient number (n>50) of patients having both CNA profile and overall survival data available for analysis. Here, the absolute activity of CNA signature was used. For mutation data, the absolute activity is explained as the estimated mutation count contributed by a signature; similarly, for copy number data, the absolute activity is explained as the estimated segment count contributed by a signature. Kaplan-Meier survival analysis was performed using the R package “survival” with log-rank test, and Cox-proportional hazard analysis was performed using the R package “ezcox”. The cutoff value in Kaplan-Meier overall survival analysis was determined by surv_cutpoint function of “survminer” package.

### CNA signature evolution analysis

The estimated cancer cell fraction (CCF) value of each segment is derived from ABSOLUTE [25] algorithm. Firstly, we sort CNA segments by their estimated CCF in a given sample. Next, we infer an evolutional trajectory of the CNA signature activities over the estimated ordering of the CNA segments. We convert the CNA segments ordering into a set of CCF cut points with overlapping subsets of segments by decreasing CCF. The absolute CNA signature activities (counts of CNA segments) of the CNA segment subset are calculated based on single sample CNA signature fitting with TCGA CNA signature set using quadratic programming.

### Statistical Analysis

Data between two groups were compared using a two-tailed unpaired Student’s t-test or Wilcoxon rank-sum test (also known as ‘Mann–Whitney’ test) depending on normality of data distribution (Typically, preprocessed expression data are normally distributed and mutation data show non-normal distribution). Correlation analysis was performed using the Spearman method. All reported *P*-values are two-tailed, and for all analyses, *P*≤0.05 is considered statistically significant, unless otherwise specified. Multiple testing *P*-values were corrected by Benjamini–Hochberg FDR method. ‘ns’ for non-significant (*P* >0.05); ‘*’ for *P*≤0.05; ‘**’ for *P*≤0.01; ‘***’ for *P*≤0.001; ‘****’ for *P*≤0.0001. All statistical analysis was performed using R v4.3.

## Data availability

PCAWG patients’ data including allele-specific copy number, tumor purity, tumor policy, tumor WGD status and general phenotype were obtained from https://pcawg.xenahubs.net [20, 34]. PCAWG gene expression data in count format were obtained from ICGC data portal (https://dcc.icgc.org/releases/PCAWG). Chromothripsis results detected by ShatterSeek were obtained from http://compbio.med.harvard.edu/chromothripsis/. PCAWG amplicons (including ecDNAs) detected by AmpliconArchitect were obtained from Kim et al. study [35, 36]. PCAWG HRD status were obtained from Nguyen et al. study [37]. PCAWG telomere contents detected by TelomereHunter were obtained from PCAWG-structural variation working group’s study [24]. PCAWG mutational signatures (including SBS, DBS and INDEL) detected by SigProfiler were obtained from Alexandrov et al. study [12]. TCGA allele-specific copy number and gene expression in count format were obtained from GDC portal (https://portal.gdc.cancer.gov/). TCGA clinical data were obtained from https://pancanatlas.xenahubs.net.

## Code availability

All code required to reproduce the analysis outlined in this manuscript are freely available at https://github.com/XSLiuLab/Pan-cancer_CNA_signature. Analyses can be read online at https://xsliulab.github.io/Pan-cancer_CNA_signature.

## Acknowledgments

We thank ShanghaiTech University High Performance Computing Public Service Platform for computing services. We thank Raymond Shuter for editing the text. We thank multi-omics facility, molecular and cell biology core facility of ShanghaiTech University for technical help.

## Contributions

ZT, SW, CW, collected the data, developed the CNA signature analysis method and performed the computational analysis. TW, XZ, WN, GW, JW, JC, KD, FC participated in critical project discussion and resources. XSL conceptualized the idea, designed, supervised the study and wrote the manuscript.

## Key points

A mechanism-agnostic method for CNA classification and signature analysis has been constructed, this method can reveal unknown patterns of CNA compared with existing methods.

This method achieves robust and consistent results in pan-cancer copy number signature analysis compared with known methods.

Pan-cancer patterns and mutational processes for CNA signatures have been revealed through our method.

Copy number signature activity consistently predict the prognosis of cancer patients.

