## Supplementary Figure for "The repertoire of copy number alteration signatures in human cancer"

### Supplementary Figure 1

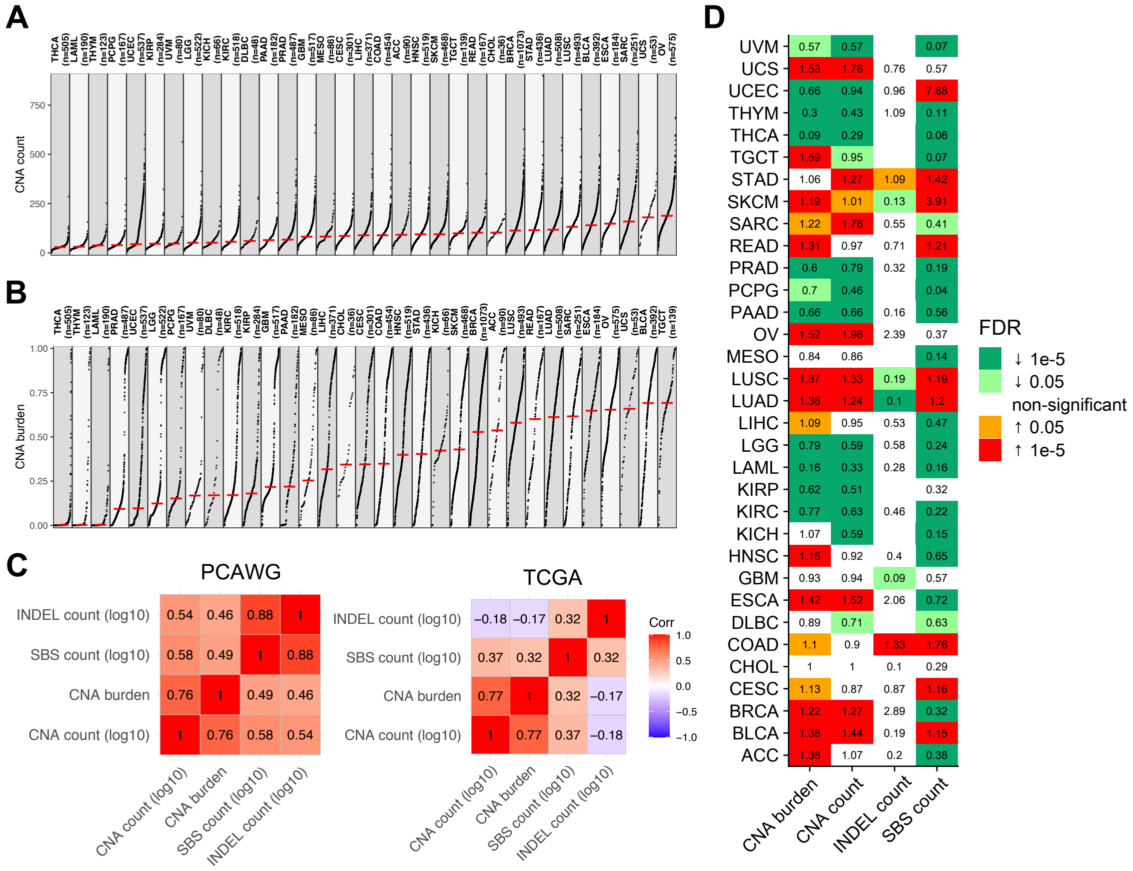


**Supplementary Figure 1. Pan-cancer distribution patterns of CNA.** (**A**,**B**) Pan-cancer distribution patterns of CNA count (**A**), CNA burden (**B**) in individual cancer types of TCGA dataset. (**C**) Correlations between CNA burden, CNA count, INDEL count and SBS count in PCAWG dataset and TCGA dataset. (**D**) Pan-cancer distribution of the enrichment scores for CNA burden, CNA count, INDEL count and SBS count in TCGA dataset. The enrichment scores are calculated as the ratio of mean value of specific cancer type compared with the mean value of the whole TCGA dataset. The colors indicate the FDR corrected *P* values of Mann-Whiney U-test.

### Supplementary Figure 2

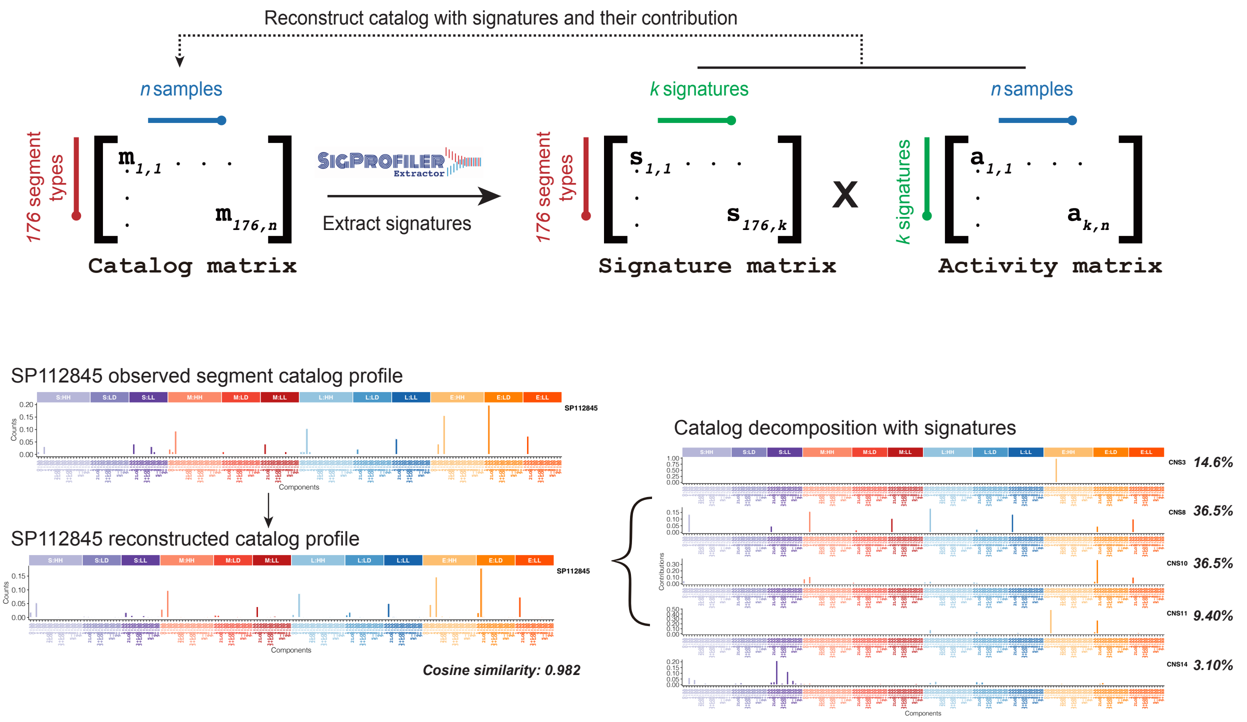


**Supplementary Figure 2. De novo CNA signature extraction with sigprofiler.**

### Supplementary Figure 3

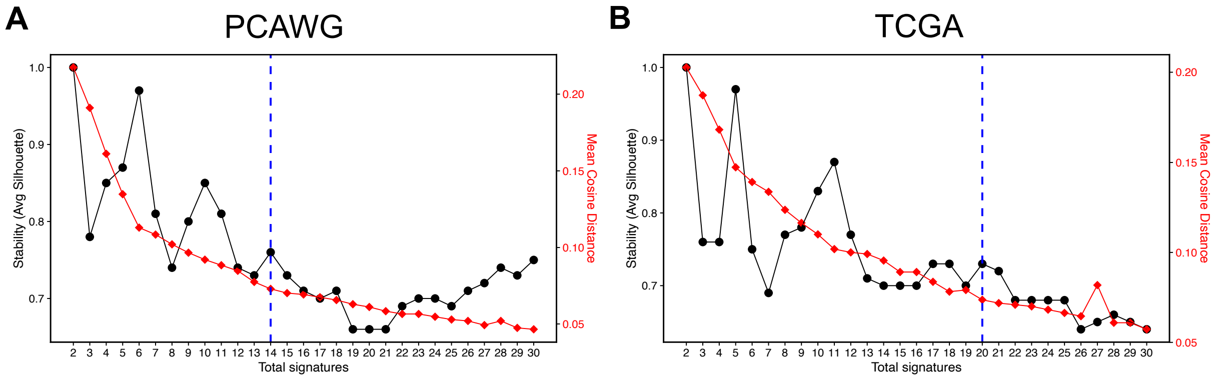


**Supplementary Figure 3. Determining the number of CNA signatures in PCAWG and TCGA dataset.** (**A,B**) The number of signatures were determined based on high reproducibility of their signatures and low error for reconstructing the original catalogs in PCAWG (**A**) and TCGA (**B**) dataset.

### Supplementary Figure 4

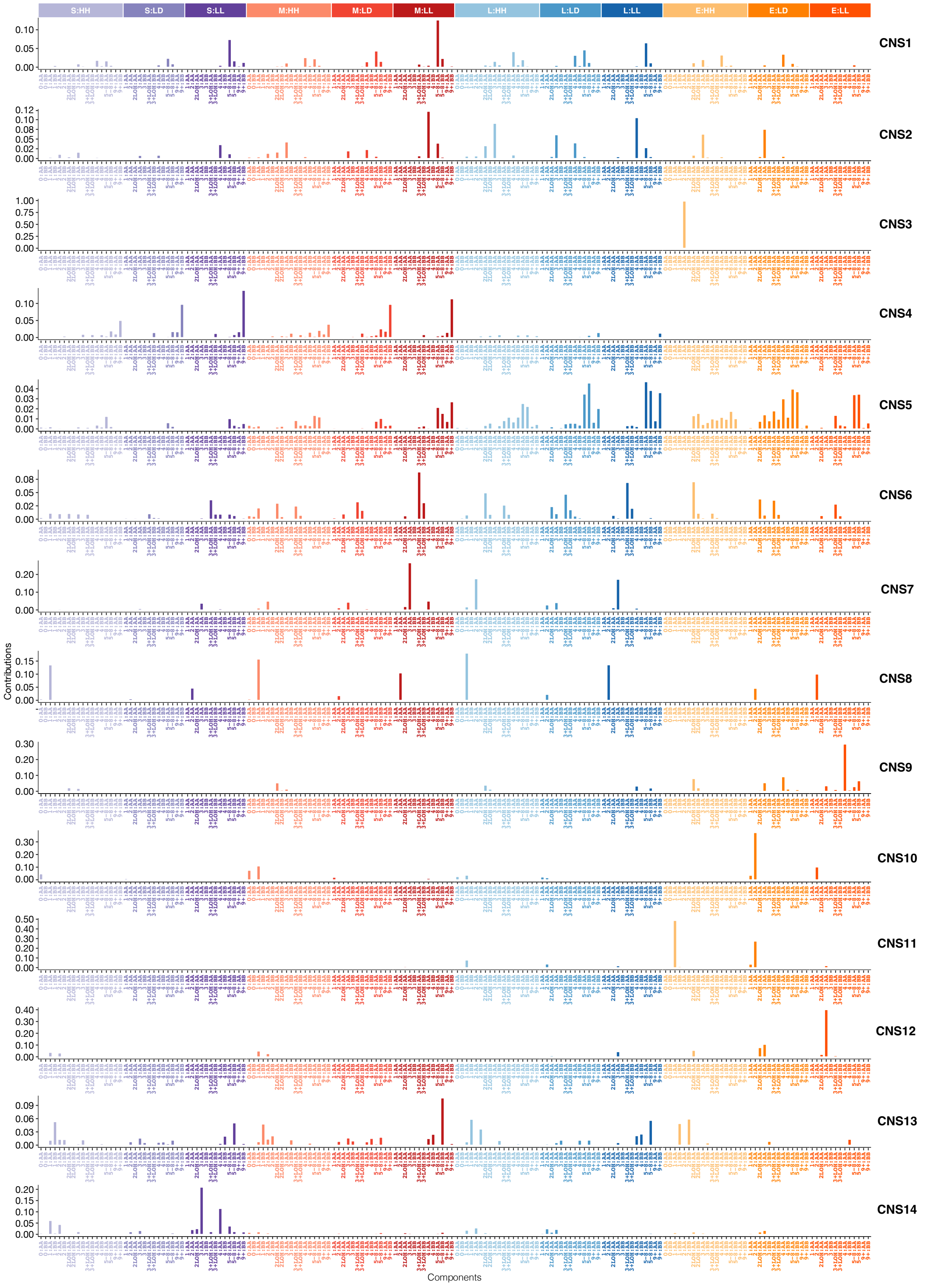


**Supplementary Figure 4. The profiles of CNA signatures extracted in PCAWG dataset.**

### Supplementary Figure 5

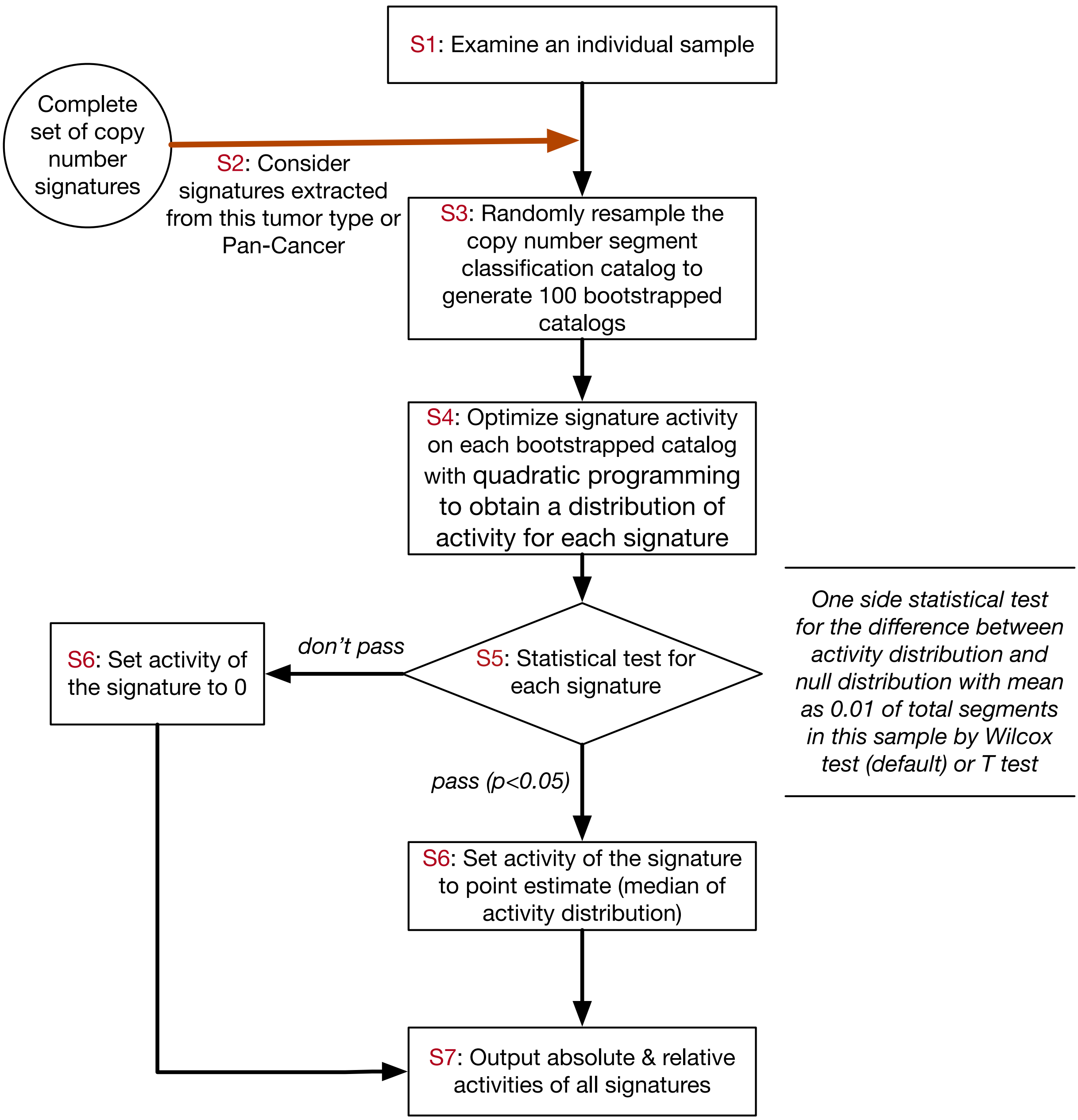


**Supplementary Figure 5. Steps for single sample CNA signature analysis.** For individual cancer patient, the CNA signatures can be identified through signature fitting process using known signatures, such as the PCAWG or TCGA CNA signatures reported in this study.

### Supplementary Figure 6

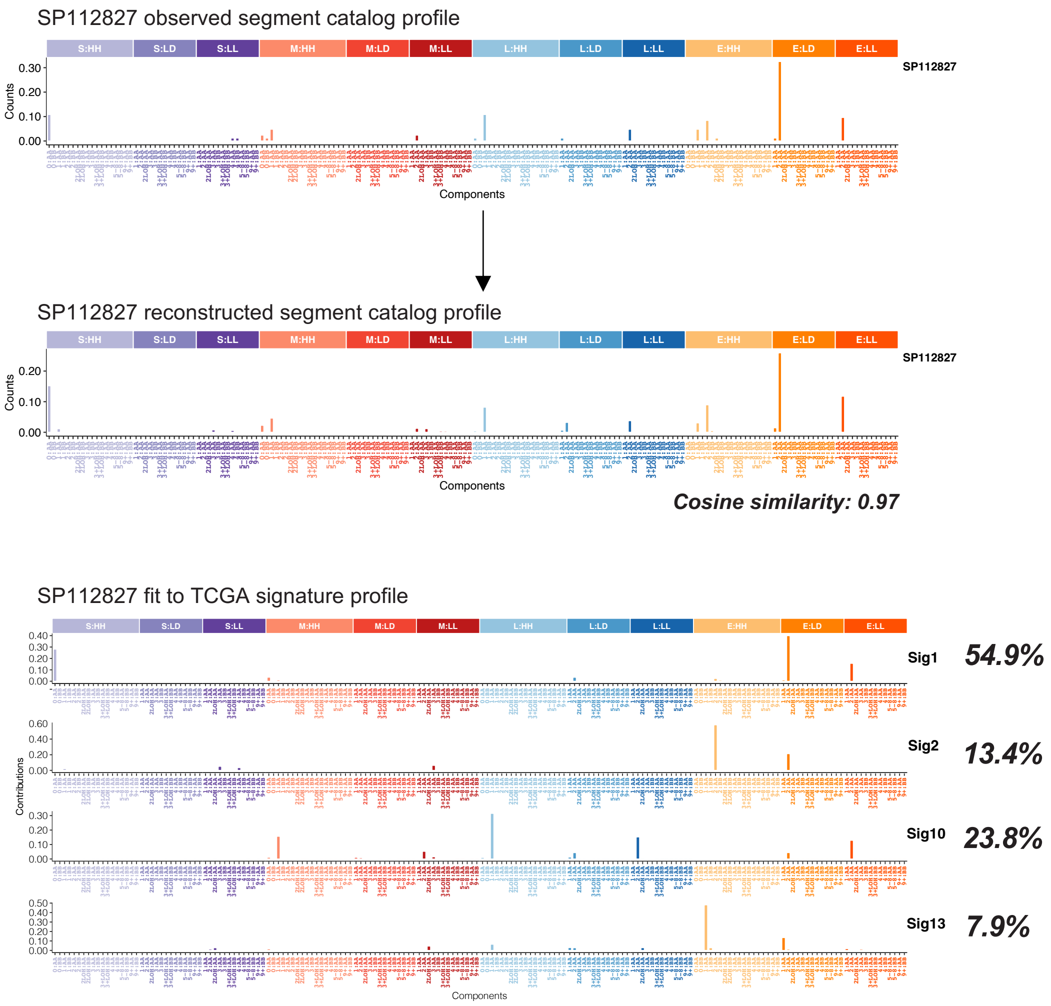


**Supplementary Figure 6. An example result of single sample CNA signature fitting.** Single sample CNA signature fitting was performed in a PCAWG sample with TCGA CNA signature set. The reconstructed segment catalog is similar to the observed segment catalogue (cosine similarity R=0.97).

### Supplementary Figure 7

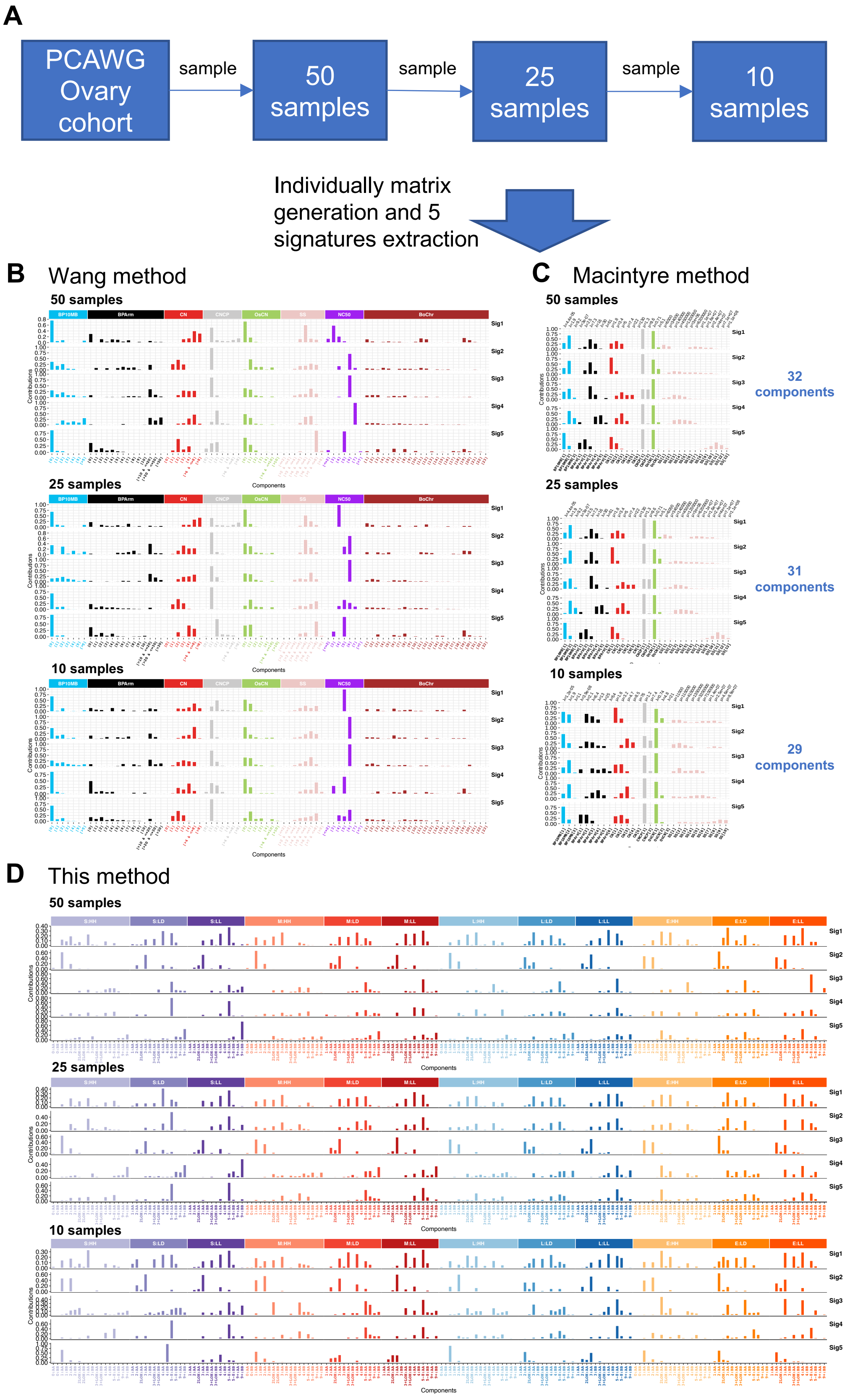

**Supplementary Figure 7. Application comparison of the three CNA signature extraction methods (Macintyre method, Wang method and the method reported here) to ovary cancer datasets with different sample size.** (**A**) The workflow of this analysis. Copy number data with different sample size are inputted for matrix generation and signature extraction using the three methods. (**B**-**D**) Signature profiles of different sample size generated through Wang method (**B**), Macintyre method (**C**) and the method reported in this study (**D**) are shown. The components of the signatures are not consistent in different datasets when using Macintyre method.

### Supplementary Figure 8

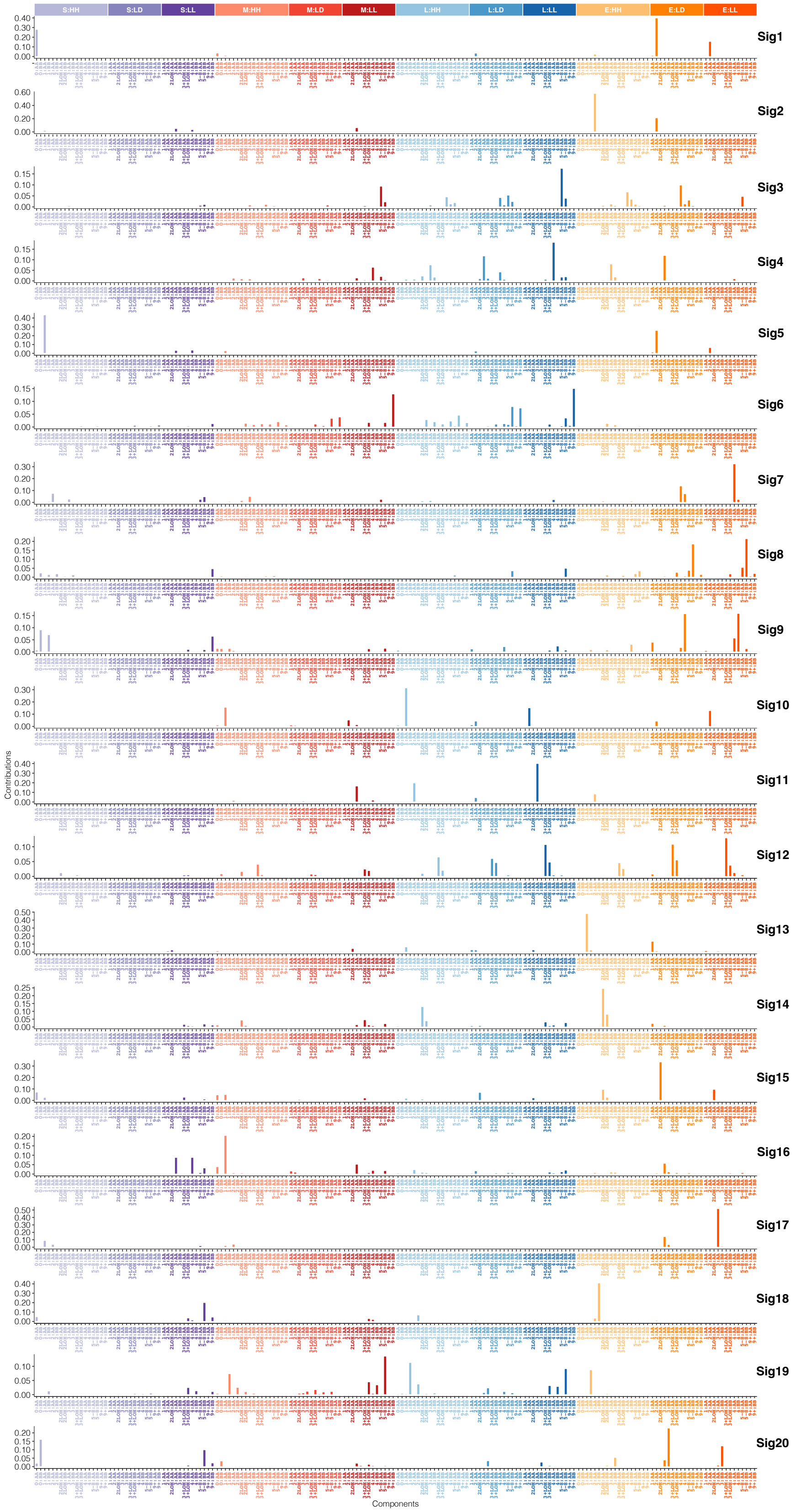


**Supplementary Figure 8. The profiles of CNA signatures extracted in TCGA dataset.**

### Supplementary Figure 9

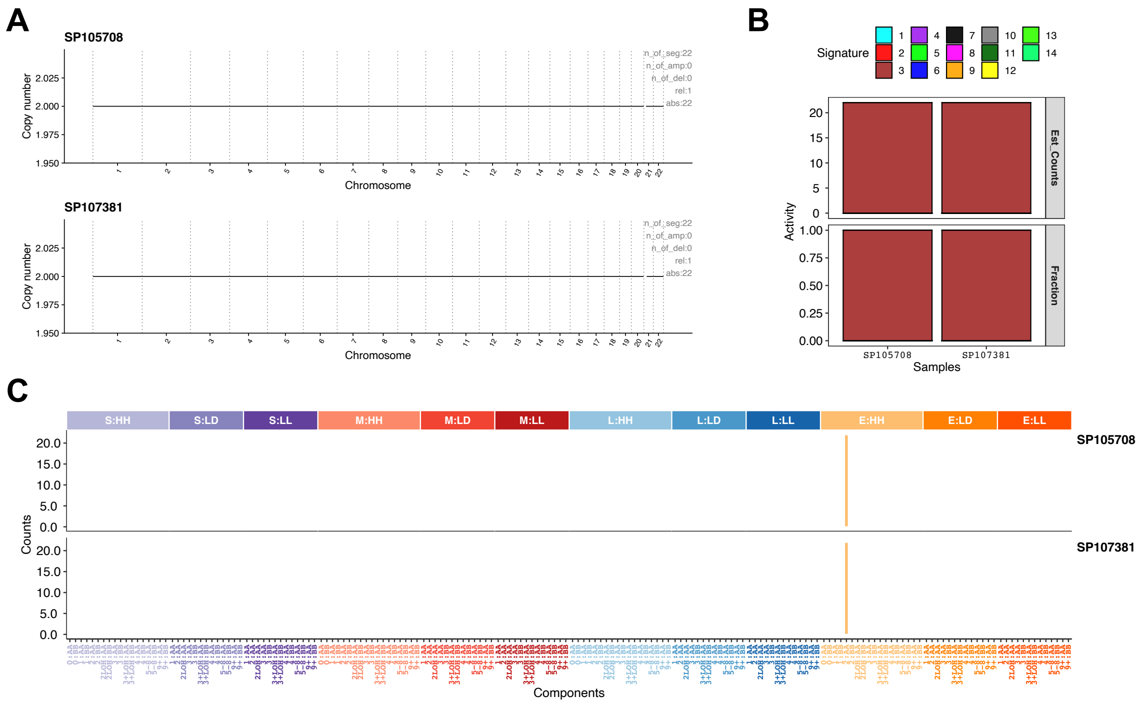


**Supplementary Figure 9. The copy number profile of samples enriched with CNS3. (A)** The copy number profile in two samples with stable genome. **(B)** CNA signature contributions of these two samples. **(C)** The CNA component profiles of these two samples.

### Supplementary Figure 10

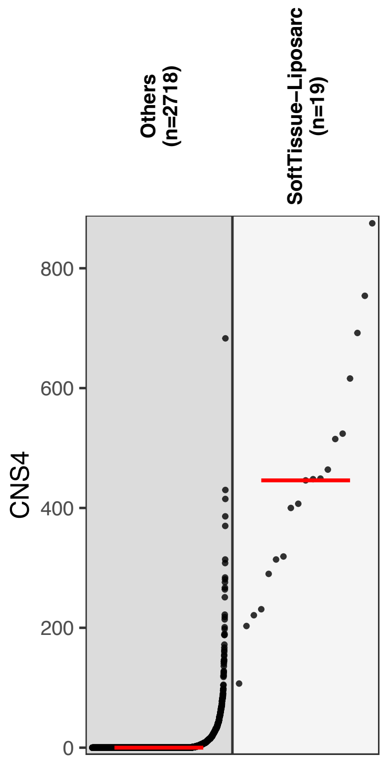


**Supplementary Figure 10. Absolute activities of CNS4 in liposarcoma and the other tumors of PCAWG dataset.** Red line indicates median CNS4 signature activity of each patient group.

### Supplementary Figure 11

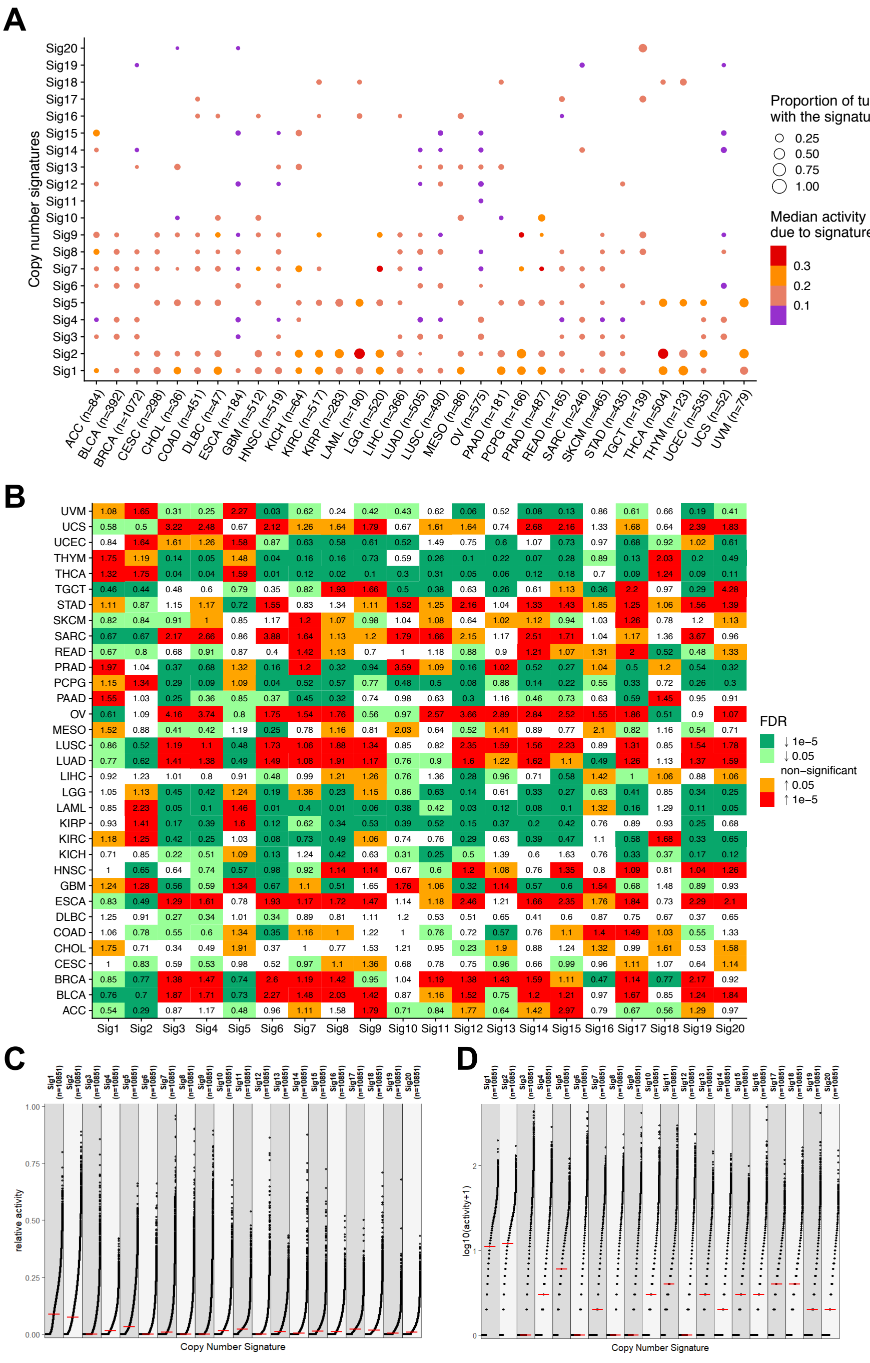

**Supplementary Figure 11. The activity distribution of CNA signatures in TCGA pan-cancer dataset.** (**A**) Proportion of tumors with the signature and the median activity of the signature are shown for 33 TCGA cancer types. For each individual tumors, only signatures that contribute ≥5% of the total are counted. (**B**) Enrichment score analysis of CNA signature in TCGA dataset. The enrichment score is calculated as the ratio of mean signature activity of specific cancer type compared with the mean signature activity of the whole PCAWG dataset. The colors indicate the FDR corrected *P* values of Mann-Whiney U-test. (**C,D**) Relative activity (percentage of the total) (**C**) and absolute activity (contributed copy number segment number) (**D**) distribution pattern of CNA signatures in TCGA dataset.

### Supplementary Figure 12

| Signatures | Representative sample profile | Prominent features | Potential mechanism |
|---|---|---|---|
| **CNS1** | 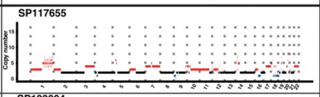  | (A) CN amplification (3-4 copies) with different size flanked by higher copy (5-8 copies) segment;<br>(B) Adjacent CN change is irregular;<br>(C) Localized amplification with various fold. | Likely Chromoanasynthesis due to MMBIR |
| **CNS2** | 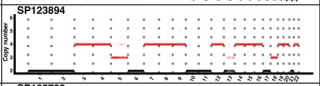  | (A) Localized CN amplification(≥3 copies <5Mb);<br>(B) Adjacent CN change is within 2;<br>(C) Oscillating CN with 3-4-3 pattern. | Unknown |
| **CNS3** | 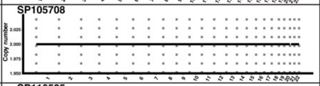  | (A) Long DNA segment (>5Mb) without copy number alteration (CN=2). | Genome without CNA |
| **CNS4** | 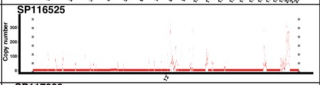  | (A) Many localized high CN amplification (copie≥9, size<5Mb);<br>(B) Adjacent CN change is >2 copies;<br>(C) Poor overall survival. | EcDNA or neochromosome |
| **CNS5** | 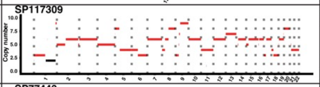  | (A) Mostly large-sized (>500Kb) DNA segment with amplification in different levels(3-8 copies);<br>(B) Adjacent CN change is irregular;<br>(C) Unequal CN amplification in the terminal of chromosome. | Likely BFB |
| **CNS6** | 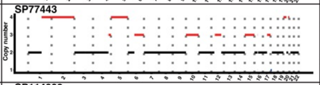  | (A) Many DNA segments of various size (>5Mb, or 50Kb-5Mb) (CN=2 or 3) with LOH;<br>(B) Adjacent CN change is irregular. | LOH and amplification |
| **CNS7** | 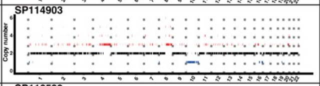  | (A) CN duplication (1-2 copies) of 50Kb-5Mb flanked by neutral segment (copy number=2)with similar size;<br>(B) Adjacent CN change is within 2 copies;<br>(C) Oscillating CN with 2-3-2 (major) and 2-3-4-3-2 (minor) pattern are obserbed. | Chromosome fragmentation and amplification |
| **CNS8** | 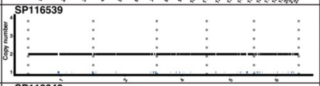  | (A) Shallow CN deletion (size<5Mb) flanked by neutral segment with similar size;<br>(B) Adjacent CN change is within 2 copies;<br>(C) Localized oscillating CN with 2-1-2 pattern. | Unknown |
| **CNS9** | 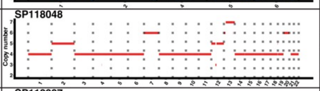  | (A) Large sized (>5Mb) CN amplification (mostly 4 copies);<br>(B) Majority of adjacent CN change is less than 2;<br>(C) The signature is associated with WGD. | WGD |
| **CNS10** | 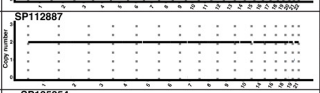  | (A) DNA segment of small to medium size(0-5Mb) (CN=0 or 1) interspersed with large sized (>5Mb) neutral segment(CN=2);<br>(B) Adjacent CN change is within 2. | Homozygous deletion |
| **CNS11** | 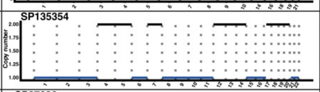  | (A) Large sized (>5Mb) shallow deletion along with larged sized (>5Mb) netural segment;<br>(B) Adjacent CN change is within 2;<br>(C) Poor overall survival. | Haploid chromosome |
| **CNS12** | 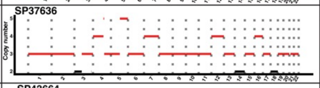  | (A) Large sized (>5Mb) DNA segment (CN=3) along with large sized (>5Mb) netural segment harboring LOH;<br>(B) Adjacent CN change is within 2;<br>(C) Poor overall survival. | Triploid chromosome |
| **CNS13** | 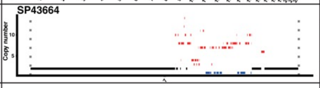  | (A) Localized high number of amplification (CN=4-8) and shallow deletions;<br>(B) Adjacent CN change is mostly >2 copies. | Unknown |
| **CNS14** | 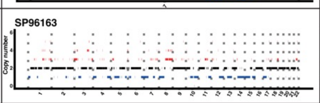  | (A) Evenly distributed small sized(<50Kb) segment (CN=1-4) flanked by DNA segment with small CN change;<br>(B) Adjacent CN change is within 2 copies;<br>(C) The signature is strongly associated with HRD. | HRD |

NOTE: To better visualize the signature features, only some chromosomes are plotted for some samples.
Abbreviations: CN: copy number; MMBIR: microhomology-mediated break induced replication; BFB: breakage-fusion-bridge; HRD: homologous recombination deficiency

**Supplementary Figure 12. Representative CNA profile, prominent features and potential mechanisms for each identified CNA signatures in PCAWG dataset.**

### Supplementary Figure 13

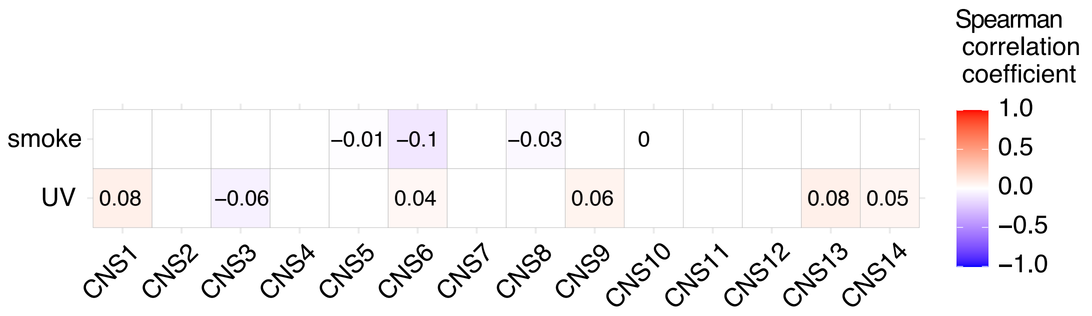


**Supplementary Figure 13. Associations between CNA signature activities and smoking or UV related SBS signatures.** Smoking related signature is a combination of COSMIC SBS4 and ID3. UV related signature is a combination of COSMIC SBS7a/b/c/d, DBS1 and ID13. Spearman correlation coefficient values are reported. UV: ultraviolet.

### Supplementary Figure 14

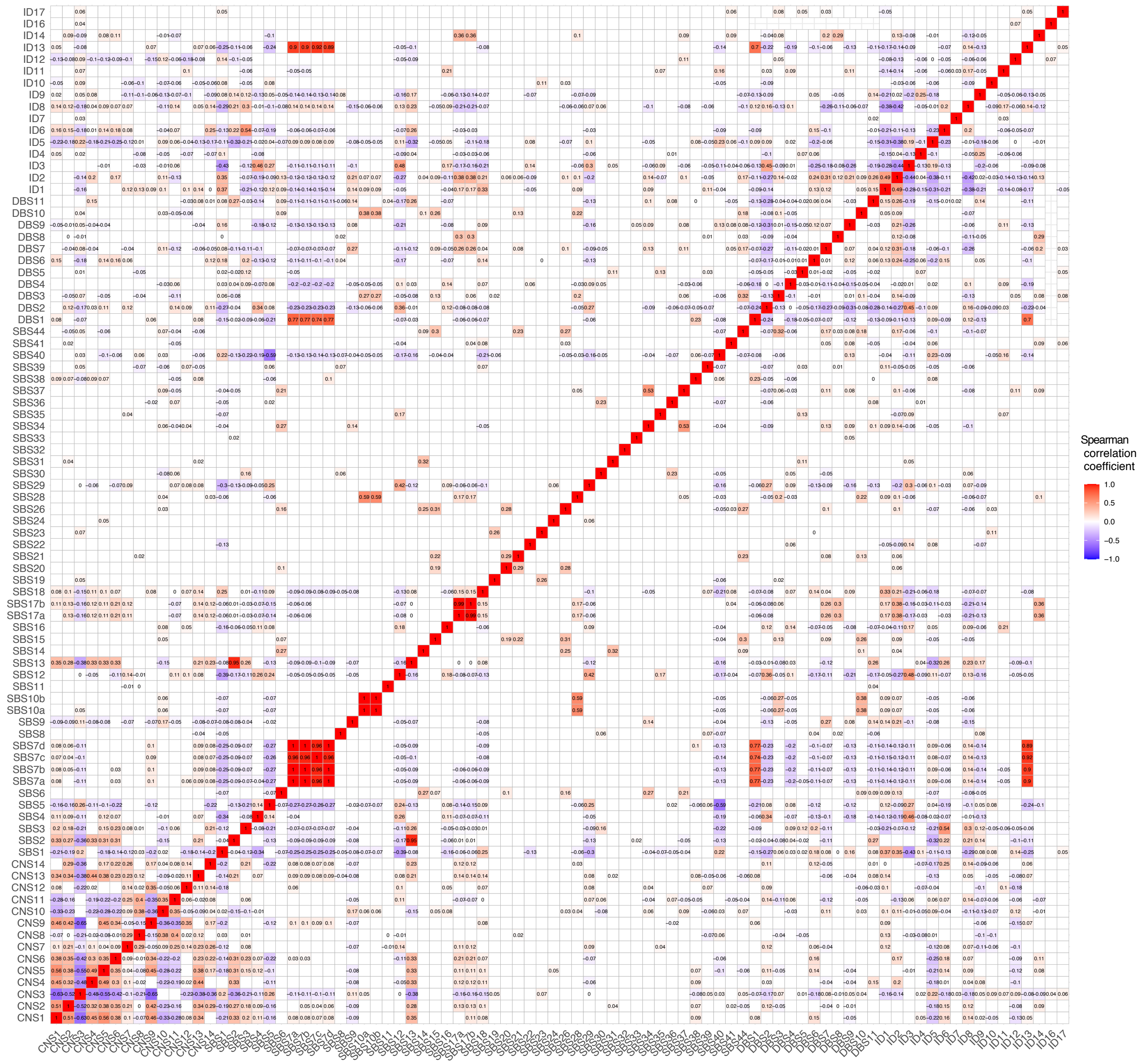


**Supplementary Figure 14. Inter-correlations between the activities of CNA and SBS signatures, DBS signatures and INDEL signatures in PCAWG dataset.** Associations between the relative activities of each signature in PCAWG dataset are calculated, Spearman correlation coefficient values are reported.

### Supplementary Figure 15

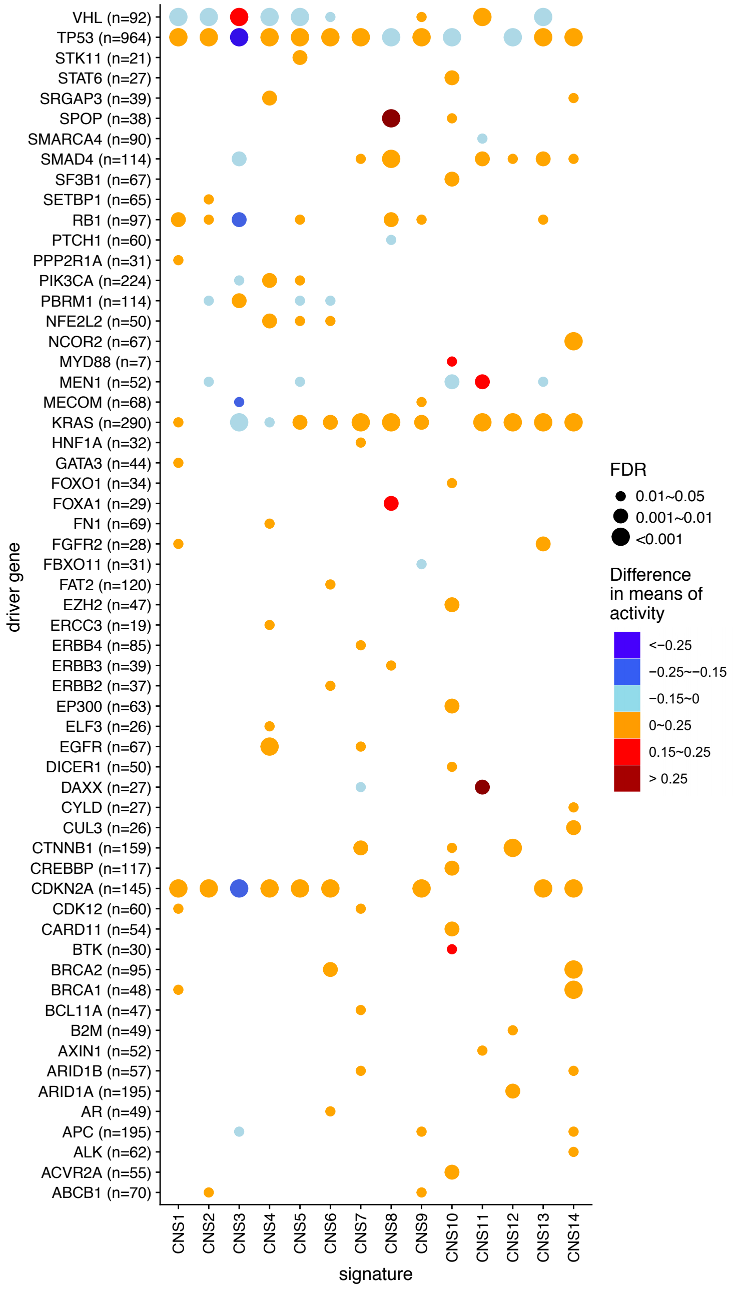


**Supplementary Figure 15. Associations between cancer driver gene mutations and the relative activities of CNA signatures.** For each cancer driver gene, PCAWG patients are divided into two groups based on the mutation status of the driver gene. The difference values in mean relative activities between two groups (activities of mutated group – activities of unmutated group) are reported, and FDR corrected *P* values (Wilcoxon rank-sum test) are shown. Initially 245 cancer driver genes are analyzed, only those genes that are significantly (FDR corrected *P* <0.05) associated with CNA signatures are shown.

### Supplementary Figure 16

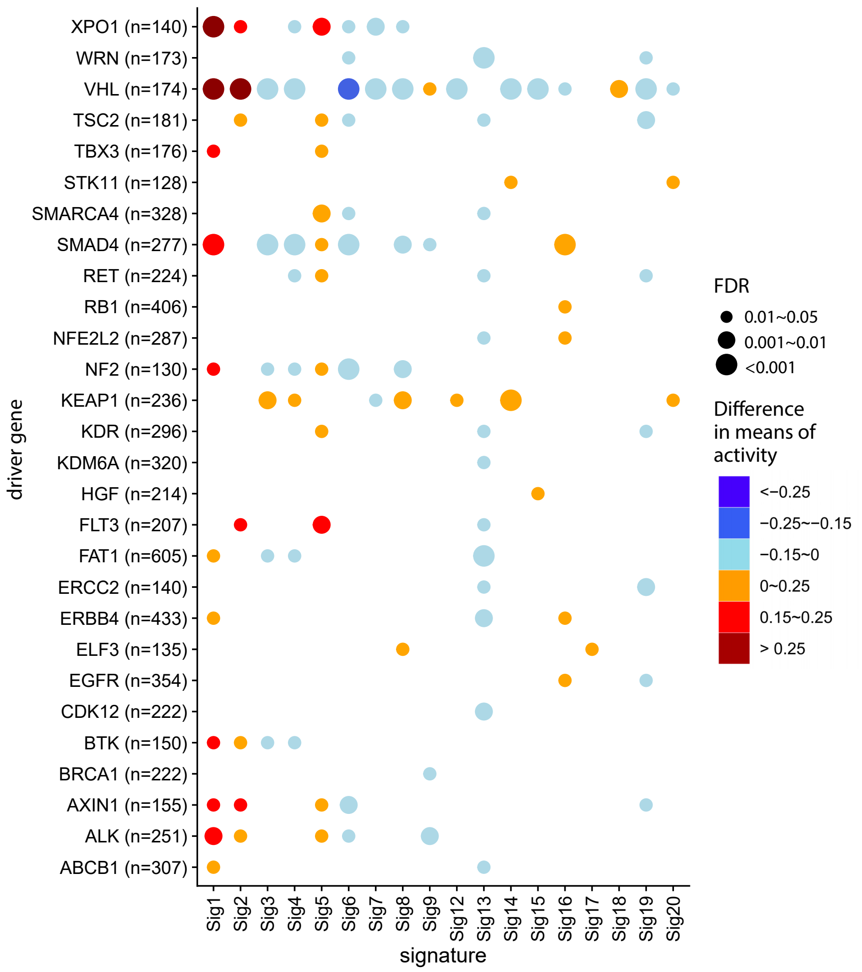


**Supplementary Figure 16. Associations between cancer driver gene mutations and the relative activities of CNA signatures in TCGA dataset.** For each cancer driver gene, TCGA patients are divided into two groups based on the mutation status of the driver gene. The difference values in mean relative activities between two groups (activities of mutated group – activities of un-mutated group) are reported, and FDR corrected *P* values (Wilcoxon rank-sum test) are shown.

### Supplementary Figure 17

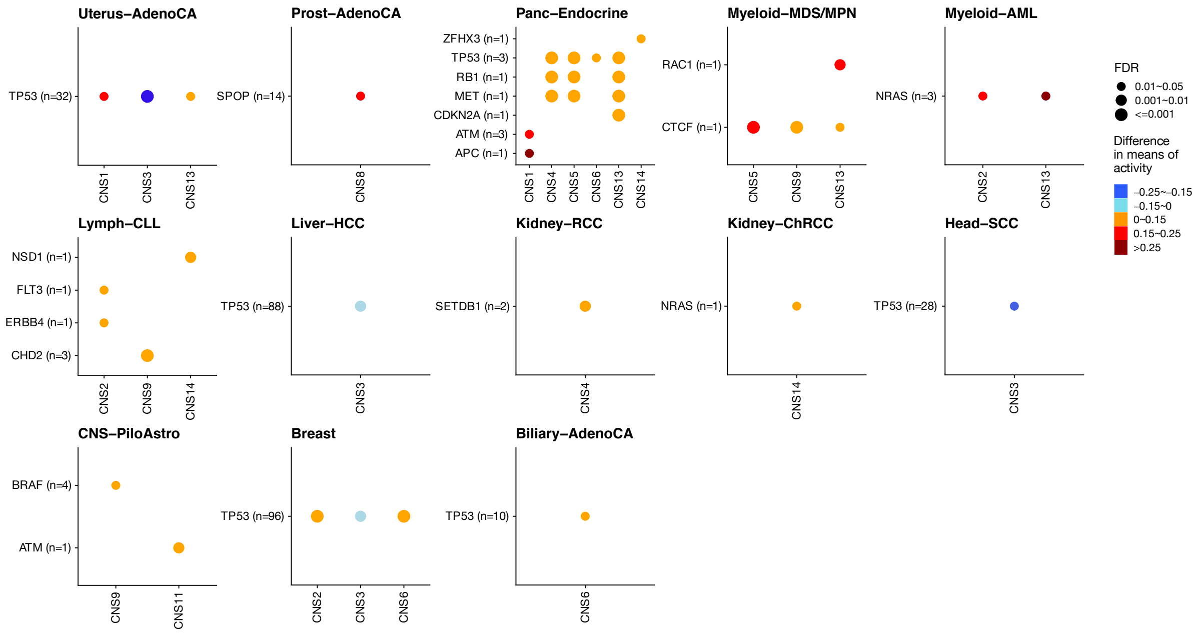


**Supplementary Figure 17. Associations between cancer driver gene mutations and the relative activities of CNA signatures in individual cancer type.** For each PCAWG cancer type, patients are divided into two groups based on the mutation status of the driver gene. The difference values in mean relative activities between two groups (activities of mutated group – activities of un-mutated group) are reported, and FDR corrected *P* values (Wilcoxon rank-sum test) are shown. Initially 245 cancer driver genes are analyzed for each cancer type, only those genes that are significantly (FDR corrected *P* <0.05) associated with CNA signatures in each specific cancer type are shown.

### Supplementary Figure 18

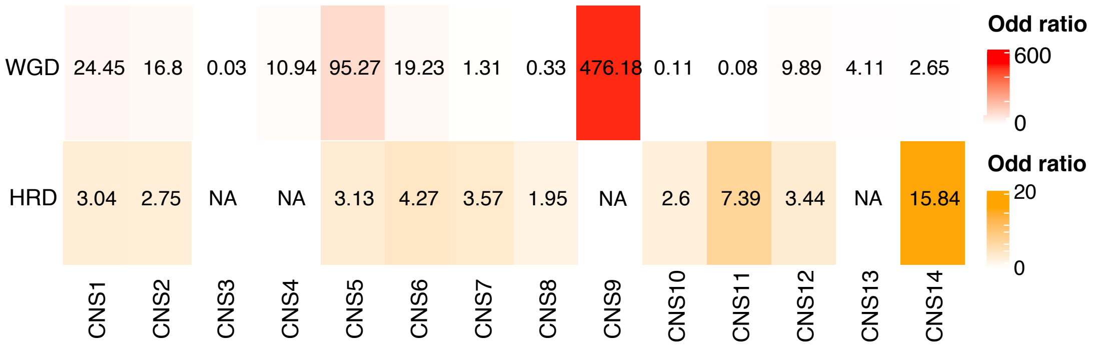


**Supplementary Figure 18. Associations between WGD, HRD status and CNA signatures.** Effect size (odds ratio) from Fisher's exact test is displayed. Significant associations with FDR adjusted *P*<0.05 are showed. CNS9 show the strongest correlation with WGD event, and CNS14 show the strongest correlation with HRD event. WGD: whole genome duplication; HRD: homologous recombination deficiency.

### Supplementary Figure 19

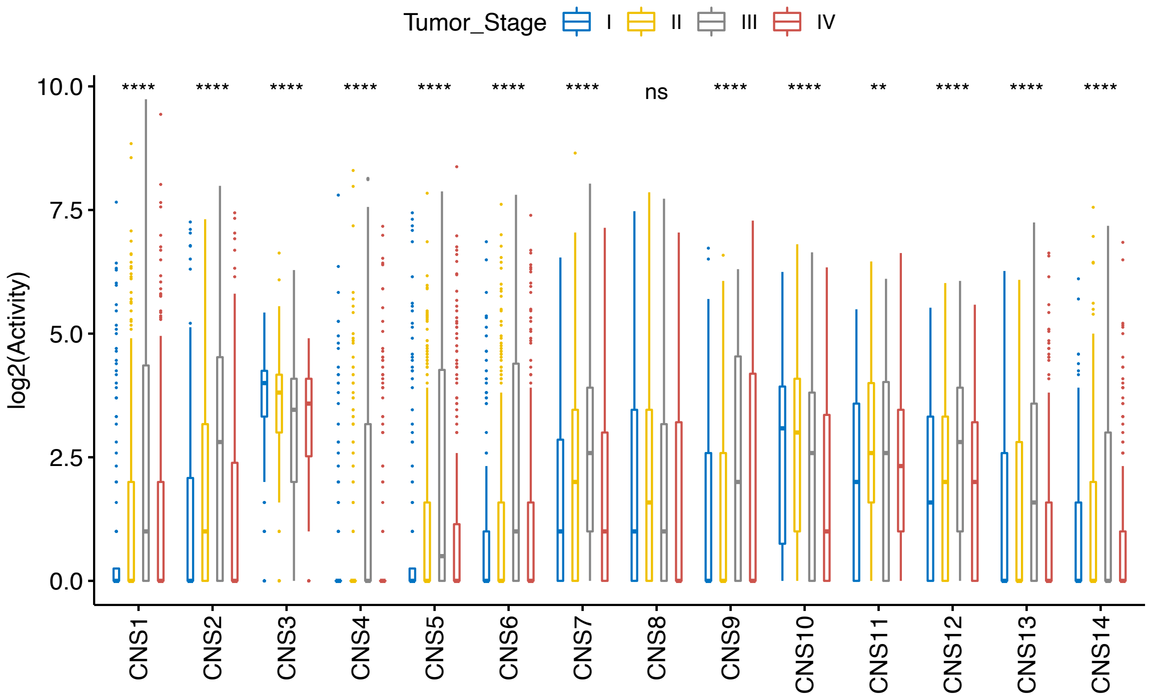


**Supplementary Figure 19. Associations between the relative activities of CNA signatures and tumor stages.** The relative activities of each CNA signature are compared in different stages of PCAWG tumors. Chi-square test p values are shown. 'ns' for *P* >0.05; '*' for *P*≤0.05; '**' for *P*≤0.01; '***' for *P*≤0.001; '****' for *P*≤0.0001.

### Supplementary Figure 20

| Signatures | Representative sample profile | Prominent features | Potential mechanism |
|---|---|---|---|
| Sig1 | 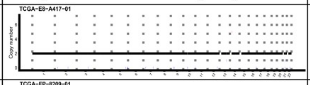 | (A) DNA segment of small to medium size(0-5Mb) (CN=0) interspersed with large sized (>5Mb) neutral segment(CN=2);<br>(B) Most of the adjacent CN changes are within 2;<br>(C) Shallow CN deletion (Size < 5Mb). | Homozygous deletion |
| Sig2 | 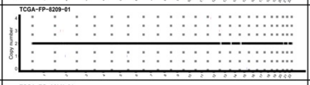 | (A) Long DNA segment ( > 5Mb) without copy number alteration (CN = 2) ; | Unknown |
| Sig3 | 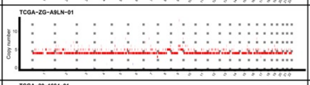 | (A) Localized CN amplification ( ≥ 3 copies <5Mb);<br>(B) Adjacent CN change is within 2;<br>(C) Oscillating CN with 4-5-4 pattern. | WGD and Chromosome fragmentation and amplification |
| Sig4 | 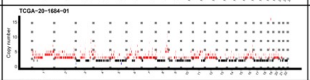 | (A) CN amplification (3-5 copies) of 50Kb-5Mb flanked by neutral segment (copy number = 4) with similar size.<br>(B) Oscillating CN with 3-4-3 (major) and 4-5-4 (minor) pattern;<br>(C) Most of the adjacent CN changes are within 2; | Chromosome fragmentation and multiple amplification |
| Sig5 | 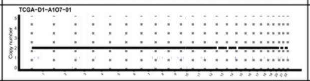 | (A) DNA segment of small to medium size(0-5Mb) (CN=1) interspersed with large sized (>5Mb) neutral segment(CN=2);<br>(B) Shallow CN deletion (size < 5Mb)<br>(C) Adjacent CN change is within 2; | Unknown |
| Sig6 | 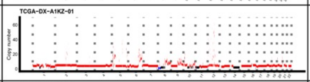 | (A) Many localized high CN amplification (copies≥9, size<5Mb);<br>(B) Adjacent CN change is irregular;<br>(C) The associated with chromothripsis. | Chromothripsis and EcDNA |
| Sig7 | 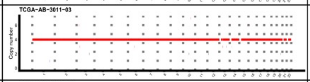 | (A) Large sized (>5Mb) CN amplification (mostly 4 copies);<br>(B) Adjacent CN change is within 2 copies;<br>(C) The signature is associated with WGD. | WGD |
| Sig8 | 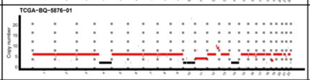 | (A) Mostly large-sized (>5Mb) DNA segment with amplification in different levels (4-6 copies);<br>(B) Adjacent CN change is within 4 copies. | Unknown |
| Sig9 | 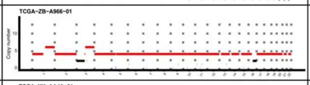 | (A) Large sized (>5Mb) CN amplification (mostly 4 copies);<br>(B) Adjacent CN change is within 2 copies;<br>(C) Oscillating CN with 4-6-4 and 4-2-4 pattern are observed. | WGD and BFB |
| Sig10 | 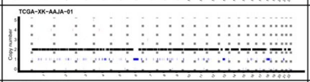 | (A) Shallow CN deletion (Size < 5Mb) flanked by neutral segment with similar size;<br>(B) Adjacent CN change is within 2 copies;<br>(C) Localized oscillating CN with 2-1-2 pattern. | Chromosome fragmentation and deletion |
| Sig11 | 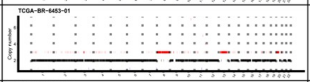 | (A) CN amplification (3 copies) of 50Kb-5Mb flanked by neutral segment (copy number = 2) with similar size.<br>(B) Most of the adjacent CN changes are within 2;<br>(C) Oscillating CN with 2-3-2 (major) pattern are observed. | Chromosome fragmentation and single amplification |
| Sig12 | 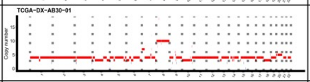 | (A) Many DNA segment of various size (>5Mb, or 50Kb -5Mb) (CN = 3 or 4) with LOH;<br>(B) Adjacent CN change is irregular. | WGD and LOH |
| Sig13 | 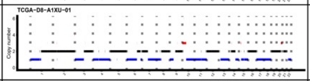 | (A) Large sized (>5Mb) shallow deletion along with large sized (>5Mb) netural segment;<br>(B) Adjacent CN change is within 2 copies. | Haploid chromosome |
| Sig14 | 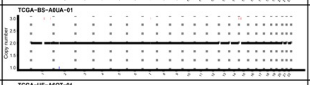 | (A) Many long DNA segment ( > 5Mb) without copy number alteration (CN = 2);<br>(B) Adjacent CN change is within 2 copies. | Unknown |
| Sig15 | 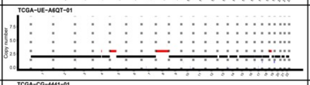 | (A) Many DNA segment of various size (>5Mb) (CN = 2 or 3) with LOH;<br>(B) Oscillating CN with 2-3-2 pattern are observed. | LOH and amplification |
| Sig16 | 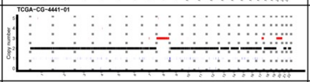 | (A) Many DNA segment of various size (>5Mb) (CN = 2 or 3) with LOH;<br>(B) Shallow CN deletion (Size < 5Mb) flanked by neutral segment with similar size; | LOH and Chromosome fragmentation and deletion |
| Sig17 | 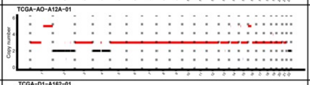 | (A) Large sized (>5Mb) DNA segment (CN = 3) along with large sized (>5Mb) netural segment harboring LOH;<br>(B) Adjacent CN change is within 2; | Triploid chromosome |
| Sig18 | 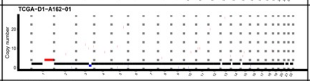 | (A) Localized high number of amplification and shallow deletions;<br>(B) Adjacent CN change is irregular. | Unknown |
| Sig19 | 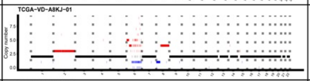 | (A) Localized high number of amplification (CN = 3-5) and shallow deletions;<br>(B) Adjacent CN change is mostly > 2 copies;<br>(C) The signature is associated with HRD. | HRD |
| Sig20 | 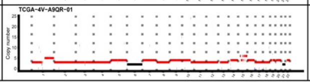 | (A) Large sized (>5Mb) CN amplification (CN = 3-5)<br>(B) Whole chromosome amplification;<br>(C) The signature is associated with TGCT. | Unknown |

**Supplementary Figure 20. Representative CNA profile, prominent features and potential mechanisms for each identified CNA signatures in TCGA dataset.**

### Supplementary Figure 21

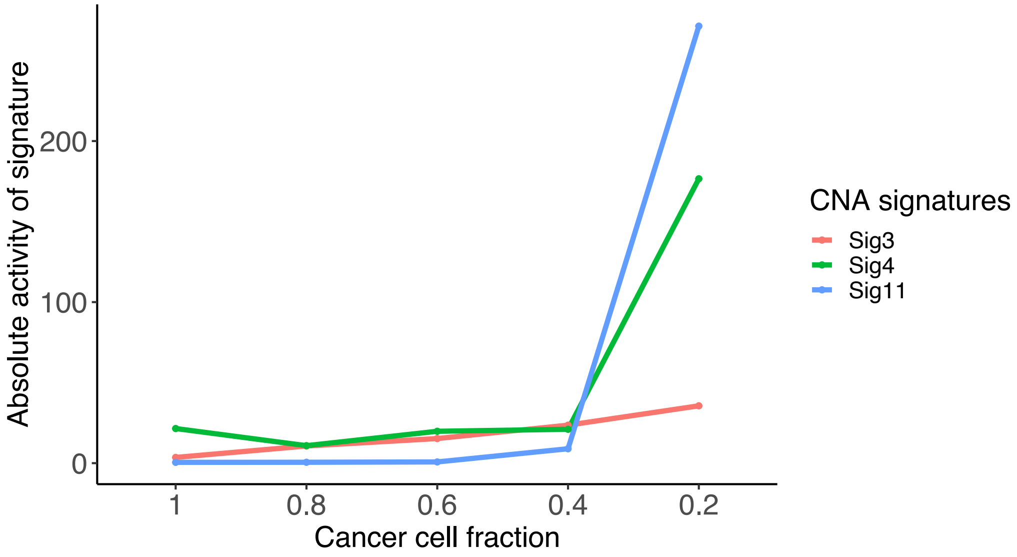


**Supplementary Figure 21. CNA signature evolution analysis.** Each line is an absolute activity trajectory (y-axis) for a single signature in one prostate cancer patient as a function of decreasing cancer cell fraction (CCF) (x-axis).

### Supplementary Table 1

| CNA classifications | Total counts | CNA classifications | Total counts | CNA classifications | Total counts |
|---|---|---|---|---|---|
| E:HH:0:AA | 90 | L:HH:5-8:BB | 1011 | M:LD:9+:BB | 3200 |
| E:HH:0:BB | 21 | L:HH:9+:AA | 105 | M:LL:1:AA | 132 |
| E:HH:1:AA | 11740 | L:HH:9+:BB | 285 | M:LL:2:AA | 2932 |
| E:HH:1:BB | 1171 | L:LD:1:AA | 638 | M:LL:2LOH:AA | 1136 |
| E:HH:2:AA | 37577 | L:LD:2:AA | 3172 | M:LL:3:AA | 8261 |
| E:HH:2:BB | 1162 | L:LD:2LOH:AA | 1444 | M:LL:3:BB | 29 |
| E:HH:2LOH:AA | 7245 | L:LD:3:AA | 4803 | M:LL:3+LOH:AA | 3214 |
| E:HH:2LOH:BB | 1544 | L:LD:3:BB | 418 | M:LL:3+LOH:BB | 1320 |
| E:HH:3:AA | 3866 | L:LD:3+LOH:AA | 1423 | M:LL:4:AA | 7258 |
| E:HH:3:BB | 588 | L:LD:3+LOH:BB | 658 | M:LL:4:BB | 508 |
| E:HH:3+LOH:AA | 571 | L:LD:4:AA | 3579 | M:LL:5-8:AA | 8322 |
| E:HH:3+LOH:BB | 334 | L:LD:4:BB | 749 | M:LL:5-8:BB | 3975 |
| E:HH:4:AA | 1987 | L:LD:5-8:AA | 3313 | M:LL:9+:AA | 747 |
| E:HH:4:BB | 475 | L:LD:5-8:BB | 2343 | M:LL:9+:BB | 4600 |
| E:HH:5-8:AA | 709 | L:LD:9+:AA | 250 | S:HH:0:AA | 1285 |
| E:HH:5-8:BB | 318 | L:LD:9+:BB | 1040 | S:HH:0:BB | 229 |
| E:HH:9+:AA | 14 | L:LL:1:AA | 98 | S:HH:1:AA | 6272 |
| E:HH:9+:BB | 20 | L:LL:2:AA | 3786 | S:HH:1:BB | 1462 |
| E:LD:1:AA | 1554 | L:LL:2LOH:AA | 851 | S:HH:2:AA | 2711 |
| E:LD:2:AA | 17896 | L:LL:3:AA | 7057 | S:HH:2:BB | 557 |
| E:LD:2LOH:AA | 3678 | L:LL:3:BB | 27 | S:HH:2LOH:AA | 1263 |
| E:LD:3:AA | 7904 | L:LL:3+LOH:AA | 2112 | S:HH:2LOH:BB | 278 |
| E:LD:3:BB | 678 | L:LL:3+LOH:BB | 656 | S:HH:3:AA | 2142 |
| E:LD:3+LOH:AA | 1534 | L:LL:4:AA | 6088 | S:HH:3:BB | 739 |
| E:LD:3+LOH:BB | 535 | L:LL:4:BB | 500 | S:HH:3+LOH:AA | 422 |
| E:LD:4:AA | 5235 | L:LL:5-8:AA | 5601 | S:HH:3+LOH:BB | 399 |
| E:LD:4:BB | 832 | L:LL:5-8:BB | 3492 | S:HH:4:AA | 1198 |
| E:LD:5-8:AA | 1560 | L:LL:9+:AA | 273 | S:HH:4:BB | 576 |
| E:LD:5-8:BB | 1365 | L:LL:9+:BB | 1479 | S:HH:5-8:AA | 1300 |
| E:LD:9+:AA | 32 | M:HH:0:AA | 2145 | S:HH:5-8:BB | 982 |
| E:LD:9+:BB | 112 | M:HH:0:BB | 317 | S:HH:9+:AA | 339 |
| E:LL:1:AA | 108 | M:HH:1:AA | 9657 | S:HH:9+:BB | 1584 |
| E:LL:2:AA | 5418 | M:HH:1:BB | 932 | S:LD:1:AA | 295 |
| E:LL:2LOH:AA | 803 | M:HH:2:AA | 3061 | S:LD:2:AA | 585 |
| E:LL:3:AA | 9780 | M:HH:2:BB | 527 | S:LD:2LOH:AA | 172 |
| E:LL:3:BB | 54 | M:HH:2LOH:AA | 3377 | S:LD:3:AA | 1213 |
| E:LL:3+LOH:AA | 1732 | M:HH:2LOH:BB | 768 | S:LD:3:BB | 188 |
| E:LL:3+LOH:BB | 368 | M:HH:3:AA | 2774 | S:LD:3+LOH:AA | 332 |
| E:LL:4:AA | 8746 | M:HH:3:BB | 787 | S:LD:3+LOH:BB | 688 |
| E:LL:4:BB | 494 | M:HH:3+LOH:AA | 1039 | S:LD:4:AA | 888 |
| E:LL:5-8:AA | 1999 | M:HH:3+LOH:BB | 685 | S:LD:4:BB | 314 |
| E:LL:5-8:BB | 2846 | M:HH:4:AA | 1621 | S:LD:5-8:AA | 1548 |
| E:LL:9+:AA | 40 | M:HH:4:BB | 876 | S:LD:5-8:BB | 1382 |
| E:LL:9+:BB | 178 | M:HH:5-8:AA | 1596 | S:LD:9+:AA | 544 |
| L:HH:0:AA | 837 | M:HH:5-8:BB | 1259 | S:LD:9+:BB | 3113 |
| L:HH:0:BB | 142 | M:HH:9+:AA | 345 | S:LL:1:AA | 103 |
| L:HH:1:AA | 9299 | M:HH:9+:BB | 1246 | S:LL:2:AA | 1638 |
| L:HH:1:BB | 1148 | M:LD:1:AA | 582 | S:LL:2LOH:AA | 546 |
| L:HH:2:AA | 6090 | M:LD:2:AA | 1143 | S:LL:3:AA | 4607 |
| L:HH:2:BB | 826 | M:LD:2LOH:AA | 657 | S:LL:3:BB | 44 |
| L:HH:2LOH:AA | 4309 | M:LD:3:AA | 2527 | S:LL:3+LOH:AA | 1317 |
| L:HH:2LOH:BB | 1260 | M:LD:3:BB | 284 | S:LL:3+LOH:BB | 775 |
| L:HH:3:AA | 4531 | M:LD:3+LOH:AA | 1006 | S:LL:4:AA | 4162 |
| L:HH:3:BB | 911 | M:LD:3+LOH:BB | 1035 | S:LL:4:BB | 477 |
| L:HH:3+LOH:AA | 942 | M:LD:4:AA | 2050 | S:LL:5-8:AA | 5115 |
| L:HH:3+LOH:BB | 618 | M:LD:4:BB | 595 | S:LL:5-8:BB | 2492 |
| L:HH:4:AA | 2473 | M:LD:5-8:AA | 2726 | S:LL:9+:AA | 746 |
| L:HH:4:BB | 928 | M:LD:5-8:BB | 2287 | S:LL:9+:BB | 5487 |
| L:HH:5-8:AA | 1641 | M:LD:9+:AA | 698 | | |

**Supplementary Table 1. List of the CNA components used for the generation of CNA signature.**
Count for each CNA component in PCAWG dataset is shown.
